# A *k*-mer-based maximum likelihood method for estimating distances of reads to genomes enables genome-wide phylogenetic placement

**DOI:** 10.1101/2025.01.20.633730

**Authors:** Ali Osman Berk Şapcı, Siavash Mirarab

## Abstract

Comparing each sequencing read in a sample to large databases of known genomes has become a fundamental tool with wide-ranging applications, including metagenomics. These comparisons can be based on read-to-genome alignment, which is relatively slow, especially if done with the high sensitivity needed to characterize queries without a close representation in the reference dataset. A more scalable alternative is assigning taxonomic labels to reads using signatures such as k-mer presence/absence. A third approach is placing reads on a reference phylogeny, which can provide a far more detailed view of the read than a single label. However, phylogenetic placement is currently only possible at scale for marker genes, constituting a small fraction of the genome. No current method is able to place all reads originating from anywhere in the genome on an ultra-large reference phylogeny. In this paper, we introduce krepp, an alignment-free k-mer-based method that enables placing reads from anywhere on the genome on an ultra-large reference phylogeny by first computing a distance from each read to every reference genome. To compute these distances and placements, krepp uses a host of algorithmic techniques, including locality-sensitive hashing to allow inexact k-mer matches, k-mer coloring graphs to map k-mers to reference genomes, maximum likelihood distance estimation, and likelihood ratio test for placement. Our experiments show that krepp is extremely scalable, improving on alignment by up to roughly 10×, computes very accurate distances that approximate those using alignments, and produces highly accurate placements. When used in the metagenomics context, the precise phylogenetic identifications provided by krepp improve our ability to compare and differentiate samples from different environments.

## Introduction

Comparing short reads of an unknown origin against an evolutionarily diverse set of reference genomes is needed in several applications, such as metagenomics [1] and decontamination detection [2]. The reference organisms are evolutionarily related, and taxonomic or phylogenetic trees can be used to model those relationships. Since the genome generating a read may not be well-represented in the reference set (e.g., with the same species or subspecies), it is often insufficient to assign a read to a reference genome; instead, we need to characterize it relative to all references. However accomplished, this goal can be thought of as having two components: Quantifying how close a read is to each reference genome and using the results to place the read in the taxonomic or phylogenetic context.

The phylogenetic tree has a higher resolution than the taxonomy. For example, the WoL-v2 [3] phylogeny has 15,246 internal nodes compared to 3,755 in its taxonomy. A phylogeny also provides interpretable branch lengths. Placing metagenomic reads on a phylogeny enables many downstream analyses [4], including sample differentiation [5] and UniFrac calculation [6], and has even outperformed *de novo* phylogeny reconstruction in such applications [7]. Moreover, the availability of ultra-large reference trees [e.g., 3, 8] has provided reliable backbone trees to use. However, placing reads on ultra-large phylogenies is challenging because it requires not just more computational resources but also intricate pipelines with many steps, such as marker detection and alignment.

Existing methods for metagenomic identification fall into two categories—those that attempt to identify every read [e.g., 9] and those that focus on a limited set of marker genes [e.g., 10]. Marker-free methods have been content with taxonomic identification (with one exception [11]) because the phylogenetic placement of any read from anywhere in the genome faces the challenge of modeling homology across the genome in addition to high computational demands. Some marker-based methods address the more ambitious goal of phylogenetic placement, often [12–18] but not always [19, 20] by aligning reads to reference alignments. These methods, however, miss out on the vast majority of reads when data are genome-wide (as opposed to amplification or targeted capture of marker genes). Thus, we can *either* use all reads but only get taxonomic labels *or* obtain phylogenetic placement but only for a small fraction of reads. This dichotomy has been a practical necessity because scalable genome-wide read placement methods do not exist (the only existing method, App-SpaM [11] does not scale). Practitioners are left with *ad hoc* solutions such as mapping reads to genomes and assigning reads to tree leaves with the best matches [21] (Fig. 1a).

**Figure 1.**
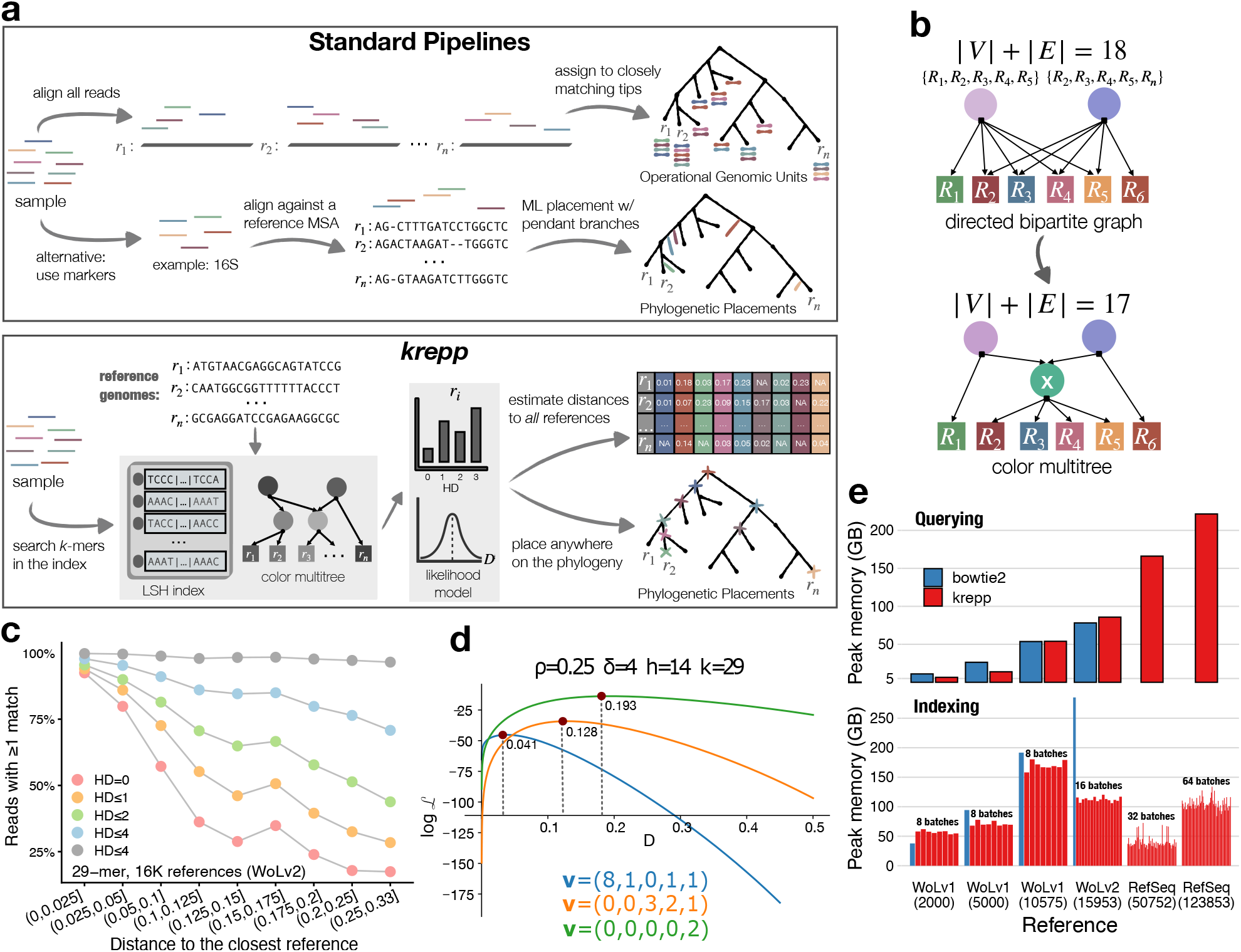
**a**, Comparison of standard pipelines, either using alignment to find operational genomic units or marker genes to perform phylogenetic placement, and krepp. **b**, Top: A trivial multitree with two non-singleton colors (circles) and six references (squares), forming a bipartite graph. Each color is simply the union of its constituents: {r1, r2, r3, r4, r5} and {r2, r3, r4, r5, r6} . Bottom: The multitree could be made smaller (in terms of total number of edges and vertices, |E| + |V|) by adding a meta-color {r2, r3, r4, r5} . **c**, As the genome-wide distance, estimated using Mash [24], between the query genome and the closest reference increases, fewer reads have any k-mers that match a reference exactly, but most have some at Hamming distance 4. **d**, The likelihood function and its maxima with a realistic parameter configuration for three different k-mer match histograms. **e**, Memory requirements of krepp, compared to bowtie2 for reference datasets with varying sizes (2000−123,853 genomes), both for indexing (bottom) and querying reads (top). krepp can build its index in batches (shown as separate bars), controlling the peak memory use.

The overarching goal of this paper is to break this dichotomy and enable genome-wide read placement. Specifically, we address two related problems. Given is a set of query reads of unknown origin and a large and evolutionary diverse set of reference genomes ℛ . i) We seek to *compute the distance* from each read q to each reference genome that is sufficiently close to q. ii) We are also given a phylogeny T leaf-labeled by ℛ and seek to *place each query read* independently onto T .

We define the read-genome distance as the Hamming distance (HD) between the source genome of q and the reference genome, restricted to the region where q is sampled from. If the rate of evolution was fixed across the genome, this would be 1 −ANI (average nucleotide identity), but since rates do vary, read-genome distances match 1 − ANI only in expectation. The best existing approach to compute such distances is aligning q to reference genomes. Efficient methods exist for aligning q to many genomes (e.g., bowtie2 [22] and minimap2 [23]), but these do not easily scale to tens of thousands of reference genomes and are less effective at high distances (> 10%).

We propose a scalable solution to both problems—a k-mer-based algorithm to compute the distance between a read and relevant reference genomes and a placement method. The resulting method, krepp (<u>k</u>-mer-based <u>r</u>ead <u>p</u>hylogenetic <u>p</u>lacement), is scalable to tens of thousands of microbial reference genomes and is accurate both in terms of distances it computes and its placements.

## Results

### krepp accurately estimates read-to-genome distances

We propose an algorithm to compute the distance between a read and a set of reference genomes. The algorithm broadly has two aspects (Fig. 1a): indexing k-mers from reference genomes in a way that allows inexact matches at the query time, and computing distance given inexact kmer matches.

#### Indexing and matching k-mers

To increase sensitivity and enable precise calculation of high read-to-genome distances, we go beyond exact k-mer search and allow query k-mers to match similar reference k-mers (measured by the Hamming distance). We achieve this by partitioning reference k-mers (default: 29-mers) into buckets using locality-sensitive hashing (LSH), a scheme adapted from the CONSULT family of methods [25– 27] with substantial changes required for distance calculations. We use the bit sampling technique to compute hash values by selecting h (default: 14) random but fixed positions from k-mers, resulting in 2^2*h*^ buckets, which together constitute the LSH index. Then, during the query time, we compute the HD between a query k-mer and all reference k-mers in the bucket specified by its hash value, and retain matches up to an HD threshold δ (default: 4) as its putatively homologous counterparts in reference genomes. Compared to exact k-mer matches, our indexing scheme boosts the portion of reads with at least one hit up to fourfold for relatively novel query genomes (>10% distance) with respect to the reference set (Fig. 1c).

After finding k-mers in the LSH index that are similar to a query k-mer, we need to identify reference genomes that include those k-mers, which is the well-studied colored k-mer representation problem [28]. Here, *color* refers to a subset of reference genomes with at least one shared k-mer (a singleton genome is also a color). The literature primarily focuses on highly similar reference genomes (e.g., pangenomes) in the context of colored de Bruijn graphs [29–31]. In our application, k-mers come from an evolutionarily diverse set, making most (but not all) colors sparse. We represent each non-singleton color as a union of other colors. This effectively defines a directed acyclic graph (DAG) G = (V, E), with colors as nodes and edges representing set partitioning. A trivial DAG makes each non-singleton color the parent of all its associated singletons (i.e., leaves of the DAG). To build a more compact DAG (w.r.t. |E| + |V|), we use the insight [32] that for similar genomes, we expect to see highly overlapping colors, which can be captured by adding “meta”-colors that represent the shared genomes (see Fig. 1b). Finding a minimal DAG has been an intractable problem and has motivated heuristic solutions such as clustering [32]. Instead, we utilize the given phylogeny and gradually build a multitree of colors using a post-order traversal, updating existing colors and adding new ones when needed at each node.

Our empirical analysis shows that our heuristic creates reasonably small multitrees with bounded height (Fig. S1). Our algorithm adds ≈500 nodes per genome for a large and diverse reference dataset WoL-v1 [33] with 10,575 microbial genomes (Fig. S1a). In addition to the size, we care about the height of colors, which has a direct impact on the running time. This consideration can conflict with the objective of finding the smallest multitree (e.g., the trivial bipartite graph has height 1). In our algorithm, the height is bounded by the phylogenetic tree T, and reference trees tend to be sufficiently balanced. On WoL-v2 [3] reference set with 15,953 prokaryotic genomes, the height of the nodes in the resulting multitree is often short, not exceeding 4 for 99.84% of k-mers (Fig. S1c). The average height of the multitree across all k-mers is 0.047, compared to 0.092 when genomes are added one by one in a random order. As a result, the inferred tree leads to more k-mers with colors of height zero (Fig. S1d). Moreover, the random addition process adds 47% more multitree nodes. The multitree of colors and the LSH index constitute the memory used by krepp. This memory consumption is comparable to or better than the standard memory-efficient alignment method, bowtie2 [22], which uses the compact FM-index [34] (Fig. 1e). Moreover, krepp requires significantly less memory when building the index due to its distributed memory algorithm, enabled by splitting the LSH index into multiple sets of buckets (i.e., batches). This approach allows creating unified indexes for ultra-large datasets exceeding 100,000 microbial genomes (a RefSeq subset [35] that we will discuss later) on standard machines with 256GB memory, whereas bowtie2 fails due to its higher peak memory usage.

#### Maximum likelihood estimation of distances

We next develop a framework to compute distances given putatively homologous matches in each reference genome for k-mers of a given read, computed using LSH index and color multitree. Considering the difficulty of modeling the complex dependencies between overlapping k-mers (which is only studied in the context of exact matches [36]) krepp ignores the position of matches and makes the simplifying assumption of independence across query k-mers. This assumption allows us to summarize homologous matches against a reference genome r as a histogram, denoted by **v**_*r*_ = (v_*r*,0_, …, v_*r*,*δ*_), where v_*r*,*d*_ is the number of matches of Hamming distance d to the reference r, and the number of mismatches u_*r*_, which is simply 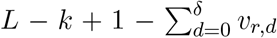, where L is the read length. For a reference r (with k-mer subsampling rate ρ_*r*_ due to minimizers), each query k-mer is either a match at a HD=d or a miss. Thus, the likelihood of having the read-wide distance D can be written as a product over all query k-mers: 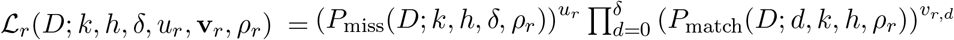 where P_match_ is the probability of observing a match of HD d and P_miss_ is the probability of no match up to the HD δ. We compute these probabilities (see Methods) using a simple substitution model and the mathematical properties of LSH. Optimizing the logarithm of this single variable likelihood function, ℒ _*r*_, results in point estimates for distances between each query read and every reference with at least one match (Fig. 1d).

#### Benchmarking accuracy of distances

We first benchmarked krepp’s maximum likelihood distances in simulations with known ground truth. We mutated 29 base genomes from Pachiadaki et al. [37] at controlled distances using a sequence evolution model with rate heterogeneity, and then generated 150bp short reads without errors. On these simulated data, krepp achieves high accuracy and very little bias in computing the true read-to-genome distance (Fig. 2a). As expected, there is noise, and the noise increases for higher distances (especially beyond 10%). The mapping rates exceed 80% even for reads with distances up to 20% and are close to perfect for less distant reads (Fig. 2b). The krepp estimates also accurately approximate the genome-wide mean and standard deviation of distances (Fig. S2). Since our simulations are simple compared to real data, we next used 15,953 real microbial genomes from WoL-v2 as reference and selected a separate set of 500 query genomes, spanning a wide range of novelty levels measured by the distance to the closest reference genome (Fig. S3b). We simulated 33 million 150bp reads across all query genomes with the default Illumina error profile, and computed read-genome distances (D) using bowtie2 run with high sensitivity and looking for all matches. We computed the genome-wide distances between each query and all reference genomes using Mash [24] and refer to it as D^*∗*^.

**Figure 2.**
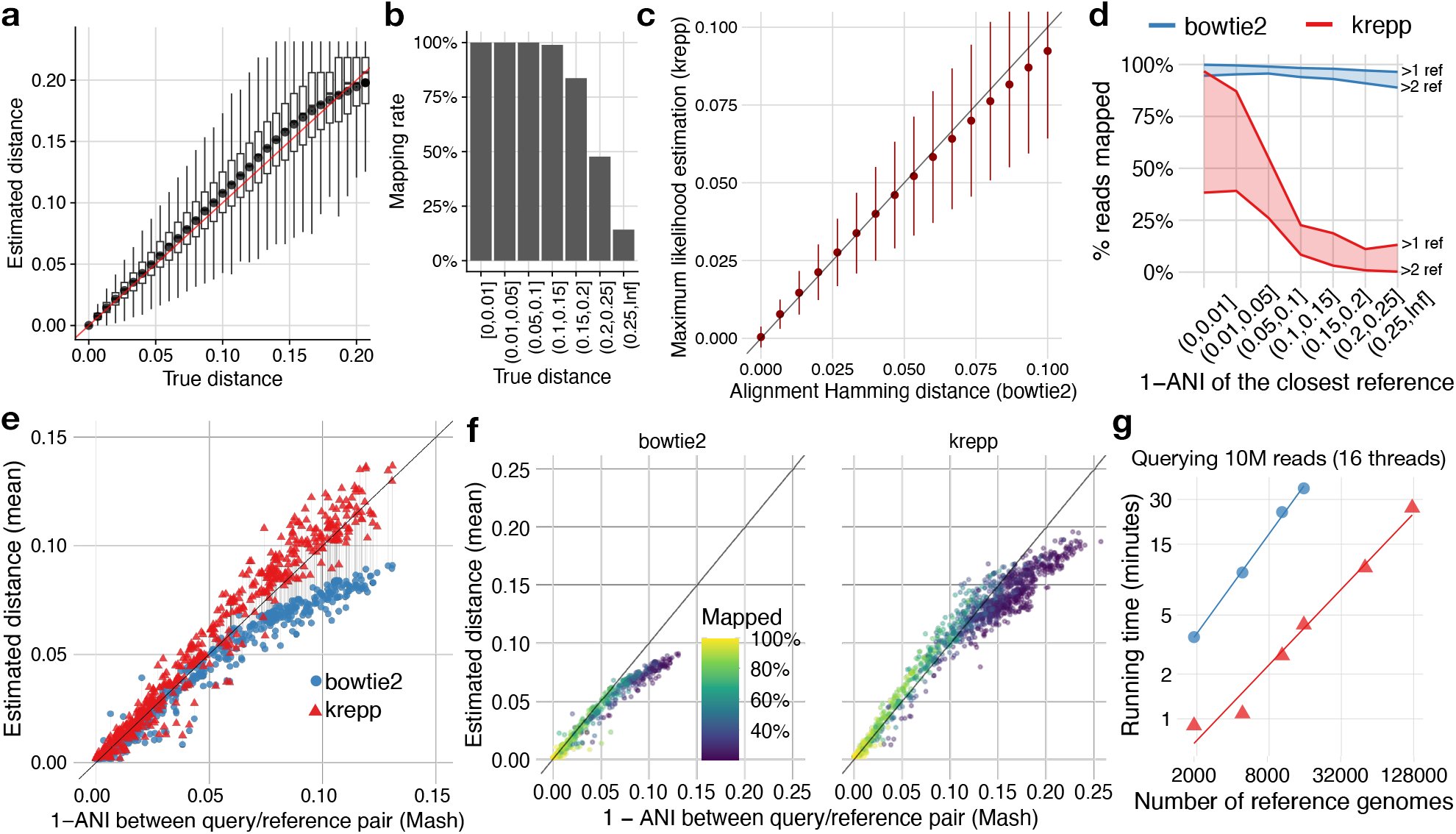
**a**, Comparing the true Hamming distance (normalized by the read length) and the estimated HD for individual 150bp short reads (1M in total). Each distribution is over 2, 500 − 50, 000 reads. **b**, Percentage of reads mapped in each true distance bin on average. **c**, ML distance estimate of krepp versus bowtie2 alignment distance, i.e., HD normalized by the read length (mean, standard deviation computed over a subsample of 1M reads). **d**, The portion of reads mapped to at least 1 (top) or 2 (bottom) references, binning queries by the genome-wide distance to their closest reference (D*′*). **e** Mean estimated distance across reads for each query/reference genome pair (connected dots) where ≥20% of reads are mapped by both methods. x-axis: D^*∗*;^ y-axis: 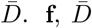 versus D^*∗*^ from each query to all WoL-v2 references with at least 20% reads mapped (colors) for each method. **g**, Running time (log-log scale) of querying 10M short reads versus the number of references. Line slops imply 𝒪 (n^0.92^) and 𝒪 (n^1.09^) growth for krepp and bowtie2. Measurements were performed on 2.25 GHz AMD EPYC 7742 CPUs using 16 threads and 256GB DRAM.

For reads with distance D ≤ 10%, where bowtie2 tends to be accurate, we take its output as ground truth. In these cases, krepp computes similar distances to bowtie2 (Fig. 2c). As in simulations, variance increases with distance, but estimates show little sign of bias. An advantage of krepp compared to alignment (e.g., using bowtie2) is that it maps more reads. More than 98% of reads are mapped to at least one and often multiple genomes using krepp, compared to 39% for bowtie2. The higher mapping rates of krepp are mostly for novel genomes (Fig. 2d). Binning query genomes by the novelty, we observe that bowtie2 fails to match 84% of the reads to *any* reference when novelty exceeds 10% whereas krepp still maps 91% of reads for these novel queries.

Notably, the high-distance matches from krepp stay accurate. Since we cannot rely on alignment results to get the ground truth for highly distant reads, to probe accuracy, we rely on genome-wide averages. We compute the mean D across all reads mapped to a reference genome 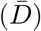. Focusing on query/reference pairs where at least 20% of reads are mapped by both methods, when D^*∗*^ < 5%, both bowtie2 and krepp are accurate, with bowtie2 performing slightly better (Fig. 2e). However, as D^*∗*^ increases to 5–12%, the bowtie2 average severely underestimates the Mash distance (1.3× on average) while krepp overestimates it to a lesser degree (1.09× on average). Examining all pairs where *each* method maps at least 20% of reads, krepp distances remain accurate on average even for 10% < D^*∗*^ < 20%, though the portion of reads mapped reduces to 50% or less (Fig. 2f). Overall, except for the most similar genomes with D^*∗*^ < 1%, bowtie2 always maps fewer reads. The mean distances from krepp more closely capture genome-wide distance D^*∗*^ than bowtie2 when D^*∗*^ > 8%, whereas, for the less novel genomes, bowtie2 has a slight advantage (Fig. S4). The range of D^*∗*^ where bowtie2 underestimates ANI coincides with the range where the efficiency of its read mapping degrades, showing that the underestimation may simply be due to unmapped reads (Fig. 2df). Computing ANI and D^*∗*^ using alternative methods such as skani [38] and orthoANI [39] shows similar trends, except for very high distances, D^*∗*^ > 15% (Fig. S5). At these levels, krepp is still more accurate in computing mean distance compared to alignment but appears to underestimate the distance computed using orthoANI and skani.

#### Scaling to ultra-large reference datasets

Beyond accuracy, krepp also enjoys better scalability than alignment with bowtie2. As the number of references increases from 2000 to 15,953 (WoL-v2), bowtie2’s running time increases 10× while krepp’s increases less than 5×, leading to >8× improvement over bowtie2 in WoL-v2 (Fig. 2g). Similarly, krepp can map 10M short reads against 50,752 (a RefSeq subset) genomes in ≈10 minutes, whereas bowtie2 can only map against 5000 (a WoL-v1 subset) genomes using the same amount of time. Both methods scale similarly with the number of reads (Fig. S6a). The total time needed to build the library is also significantly shorter for krepp than bowtie2 (e.g., 0.5× for WoLv1), and the latency of krepp can be reduced 16× by simply dividing into 16 parallel batches (Fig. S6b). Note that bowtie2 solves a more difficult problem (alignment) than krepp; these comparisons are to clarify that if the only goal is to compute the distance of a read to reference genomes, then using krepp, and hence avoiding the difficult alignment step, is preferable to bowtie2.

### krepp accurately places reads on a large phylogenies using distances

We explore several methods to use distances from reads to genomes, noting that a read can have hundreds of reference hits at low distances when the reference database is densely sampled. A commonly used approach with alignments with multiple hits is using all genomes with sufficiently small distances, which is how the operational genomic units (OGUs) are defined [21]. These OGUs can then be summarized into taxonomic profiles or assigned to tree leaves (e.g., the method Woltka). We additionally seek to find the best placement of a read on a given tree T (including on internal nodes), considering all of its read-to-genome distances. A simple baseline is placing a query q as a sister to the leaf with the minimum distance, but this approach never places q as sister to a clade, which makes the tree T moot and can easily misplace a query (Fig. S7a). For the case of ultrametric trees, a principled approach is to place q as the sister to the largest clade where all leaves have distances that are sufficiently close to the minimum distance (Fig. S7a). We propose an algorithm that follows this strategy, which can be expected to perform well if deviations of T from ultrametricity are limited. Our algorithm recursively defines distances from reads to clades in a bottom-up fashion by summarizing k-mer histograms at higher levels (see Methods). Employing a likelihood-ratio test (i.e., the χ^2^ test with 1 degree of freedom) to account for noise in estimates, we find all clades, including leaves, that are statistically indistinguishable from the minimum distance of q to any node. Among all such clades, we place as a sister to the largest, breaking ties by choosing the one with the smallest distance and leaving q unplaced if the chosen clade is the root or there are too few k-mer matches (see Methods).

We first used the WoL-v2 dataset to compare krepp placements to placing as the sister to the closest reference using distances from either krepp or bowtie2. We selected 110 query genomes from the WoL-v2 reference tree, ensuring a range of novelty levels, measured as path length on T to the closest leaf (Fig. S3). We pruned each query genome from the backbone in a leave-one-out manner, generated synthetic 150bp Illumina reads, and placed reads on the tree. We measured error as the number of edges between each placement and the original position before pruning.

The mean placement error of krepp is low despite placing most reads (Fig. 3a). Even for the most novel query genomes, the mean placement error is 3.2 edges (median: 1 edge) despite placing >50% of reads. As novelty levels decrease, more reads are placed, reaching close to 100% for novelty < 0.5. The placement error of krepp increases substantially at the highest levels of novelty (e.g., > 4). Similarly, for the least novel queries that are near identical to some references (e.g., < 0.02 novelty), the placement error is high (Fig. S7b), perhaps because placing among highly similar genomes (short branches) is challenging. Simply using the closest krepp hit increases the error. Similarly, placing each query as the sister to the lowest common ancestor of all references that are indistinguishable from the minimum krepp distance performs worse than krepp (Fig. S7c). Compared to bowtie2-closest, krepp-closest places slightly more reads and has a substantially lower errors, except for most novel queries, where it maps far more reads but has worse accuracy. If we consider unplaced reads as an error of infinity, krepp retains a median error of 0–2 in all levels of novelty. It also finds the correct edge for 26–61% of reads, depending on the novelty; bowtie2-closest has a median error of 1–4 and does not place 50% of reads across the two most novel bins (Fig. 3b). We next compared krepp to the only alternative method that can place genome-wide short reads, App-SpaM. Due to the high memory demand of App-SpaM, we restricted reference phylogenies to small clades selected from the WoL-v1 tree. For eight clades (six genera and two families, detailed in Table S2), we selected 40 or 50 genomes as references and the rest as queries (leave-all-out), which were all removed from the tree and used to simulate Illumina reads. Here, krepp is slightly more accurate and far less memory-intensive than App-SpaM (Fig. 3c). App-SpaM requires 84–189GB of memory for these small clades, and we could not build a library for two of the six clades. krepp requires only 1–6GB and is roughly 2–4x faster, including for index construction time (Fig. 3c). Both methods have similar errors, though on average, krepp has 0.15 fewer errors and places 6% more reads on the correct branch. Substantial errors by both methods underscore the difficulty of placing reads on densely sampled groups of very similar genomes. Note that App-SpaM was at least as accurate as other k-mer-based methods such as RAPPAS [20] and EPIK [19], and the alignment-based APPLES for marker-based placement [11, 19].

**Figure 3.**
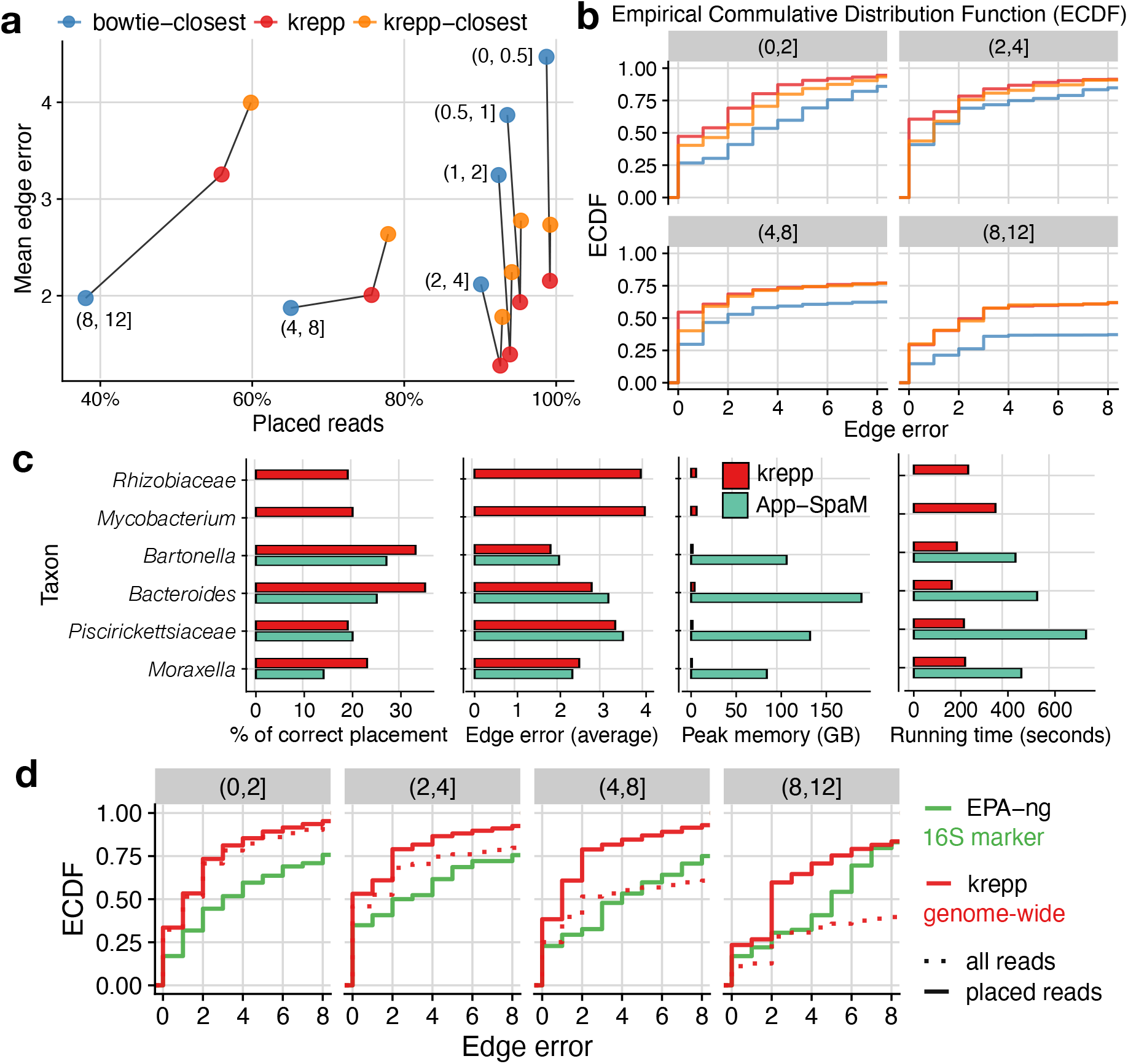
**a**,**b**, Placement error and effectiveness for 110 query genomes, binned based on novelty (measured as the path length to the closest leaf on the WoL-v2 tree), labels in **a** and titles in **b** and **d**. In **a**, unplaced reads are ignored in computing mean error; in **b**, shown error distributions treat unplaced reads as infinity error. **c**, Comparison of krepp and App-SpaM on six taxa pruned from the WoL-v1 trees. Measurements were performed on 2.10 GHz Intel Xeon Silver 4110 CPUs using 16 threads and 188GB DRAM. App-SpaM ran out of memory for *Mycobacterium* and *Rhizobiaceae*. For comparability with App-Spam, we forced krepp to place every read (i.e., including root placements and no filtering based on the number of matches). **d**, Comparing 16S marker-based placement using maximum likelihood method EPA-ng to genome-wide read placement using krepp on 100 query genomes. We show the error distribution for placed reads separately (dashed versus solid) since EPA-ng places all reads, but is limited to 16S rRNA genes, and the total number of reads analyzed differs immensely (608 versus 1M).

Finally, we ask a broader question: Can the marker-free method krepp be competitive or better than using marker-based placement using aligned reads? To test this, we selected 100 genomes that included the 16S marker gene as queries and removed them from WoL-v1 (10,575 tips) and used the rest as the reference for krepp. We generated 150bp reads from 16S marker genes of query genomes,

which, unlike genome-wide reads used for krepp, were *error-free*. We aligned these reads using WITCH [40] and placed them using the maximum likelihood method EPA-ng on the reference tree induced to species that include at least one 16S. The edge error of genome-wide reads using krepp is substantially lower than 16S reads using EPA-ng—2.36 versus 5.5 edges on average (Fig. 3d). While EPA-ng places all 16S marker reads, placement rates of krepp depend on the novelty, ranging from close to 100% for the least novel bins to 20% for those with novelty above 10 substitutions per-site (Fig. S7d). Regardless, since we have far more genome-wide reads than 16S, the total number of genome-wide reads placed remains three orders of magnitude higher for krepp compared to reads from 16S genes for EPA-ng (608 versus 1M).

### krepp improves metagenomic sample differentiation

To apply krepp on real metagenomic data, we first re-analyzed a subset of The Human Microbiome Project (HMP) [41] data, consisting of 210 samples, each with 1M subsampled paired-end short Illumina reads [21]. These samples represent seven body sites from both male (n =138) and female (n =72) subjects. We compared krepp’s ability to characterize these samples against the OGU approach of Woltka [21] (heavily reliant on bowtie2) and taxonomic profiles estimated by Bracken [42]. For krepp, in addition to placements, we also computed OGUs by assigning a read to all leaves of the tree with statistically indistinguishable distances to the minimum. We computed pairwise distance between samples using weighted UniFrac for phylogenetic profiles (krepp and Woltka) or Bray-Curtis for taxonomy-based profiles (Bracken). We measured the agreement of these distances and the available body part labels using PERMANOVA pseudo-F statistics [43].

Visualized using PCoA, samples separate well in three dimensions using krepp (Fig. 4a). Quantified by pseudo-F, krepp resulted in slightly better separation of body sites compared to Woltka (Fig. 4b), noting that OGUs of krepp are better than OGUs of bowtie-2, and placements of krepp are better than its OGUs. Here, the vast majority of reads that are mapped by krepp are also placed (Fig. 4c). Bracken is far less effective than OGUs and placements, a pattern that Zhu et al. [21] attributed to better resolution of phylogeny-based sample distances compared to taxonomy [21].

**Figure 4.**
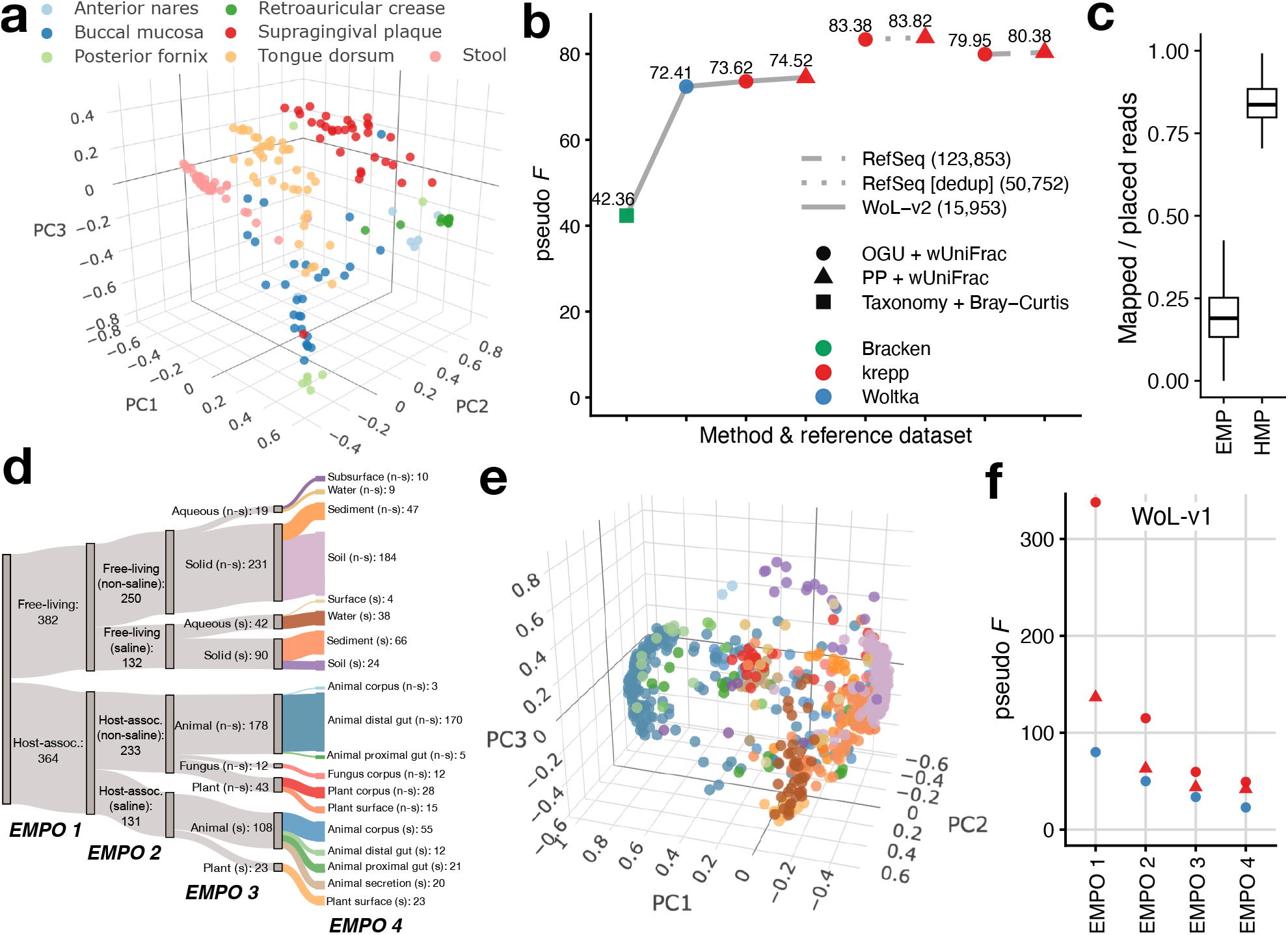
**a, d**, PCoA of weighted UniFrac distances across samples from Human Microbiome Project (n = 210) and Earth Microbiome Project (n = 746) computed based on krepp’s placements using WoL-v2 and WoL-v1 reference datasets, colored by body sites and microbial environments (EMPO 4), respectively. **c**, Ratio of total counts in BIOM tables obtained before rarefication using OGU approach krepp’s ditances (i.e., mapped) and krepp’s placements, per sample, in Earth’s microbiome (EMP) and human microbiome (HMP) data. **d**, Separation of body sites, quantified by pseudo-F statistic computed on distance matrices obtained using different methods and reference datasets (WoL-v2, RefSeq before and after deduplication). **e**, Organization of microbial environments according to Earth Microbiome Project Ontology categories. For EMPO 3 and EMPO 4, n-s: non-saline; s: saline. **f**, Separation of microbial environments at different levels, quantified by pseudo-F statistic using WoL-v1 reference dataset.

The real power of krepp for human microbiome analysis is enabling much larger reference data sets. Unlike the alignment-based Woltka, krepp is capable of indexing much larger datasets, enabling us to move beyond WoL references to build indices with 123,853 and 50,752 microbial genomes. The difference between the two sets is that the smaller one removes near-duplicate genomes. While krepp provides small improvements in sample differentiation when WoL-v2 is used, the pseudo-F statistic of krepp substantially increases with the larger reference trees (Fig. 4b). These improvements are likely due to better representation of relevant species. Surprisingly, the larger tree with duplicates included does not attain better separation of body sites compared to the deduplicated tree. The reduced separation can be partially explained by the increased ability to capture variation within body-sites. For instance, we observed higher distances within the stool category, but anterior nares to supragingival plaque distances decrease by 4% on average (Fig. S8). Regardless, with both of these trees, krepp’s placements enjoyed higher pseudo-F statistics compared to the OGU approach.

Since the human microbiome is relatively well-studied, we next analyzed more novel microbial communities where queries often lack sufficient representation in reference sets [37, 44]. For this, we used Earth Microbiome Project [45], which includes a wide variety of samples from non-human hosts and free-living in environments such as soil and water. We analyzed a subset consisting of 746 shotgun metagenomic samples, each with roughly 2M reads on average (in total 14.7B), that were selected and analyzed using Woltka (WoL-v1 reference dataset) by Shaffer et al. [46]. Earth Microbiome Project Ontology (EMPO version 2) organizes these samples into categories at four levels (Fig. 4d) based on host association (EMPO 1), salinity (EMPO 2), host taxon (for host-associated) or phase (for free-living) (EMPO 3), and the specific environment (EMPO 4). We used the same analyses as HMP datasets, focusing on comparing krepp to Woltka at different ontology levels, using the same reference set (WoL-v1) as used by Shaffer et al. [46].

On the EMP data, krepp continues to separate samples by their environmental properties (Fig. 4e). Measured again using pseudo-F statistic, the distance matrices computed from krepp OGUs reflect EMPO categories remarkably better than alignment-based OGUs at all four levels (Fig. 4f). While improvements are observed in every ontological level (4.2×, 2.3×, 1.8×, and 2.1× higher at EMPO 1–4, resp.), they are larger at higher levels. Applying the pairwise PERMANOVA test to 19 environment categories at EMPO 4, we observe that 87% of pairs show significant differences (Fig. S9). Moreover, cases of insignificant differentiation are highly overrepresented among pairs that include surface (non-saline) and animal corpus (non-saline), which included only four and three samples, resp. Conversely, soil categories are highly distinguished from all other environments, including between saline and non-saline categories.

While krepp’s placements lead to substantially higher pseudo-F statistics than alignment-based OGUs from Woltka, they are not better than krepp’s OGUs (Fig. 4f). The cause of this pattern, which is in contrast to HMP results (Fig. 4b), is not fully clear but may be related to the treatment of ambiguous read mappings during placement. When there are numerous reference hits in divergent clades with distances that are high (e.g., >15%) but indistinguishable, the OGU approach still counts each hit with a weight disproportional to the number of such hits. However, such reads will most often be placed higher on the tree, having little impact on the weighted UniFrac distance compared to the OGU approach. Consistent with this explanation, on EMP, many reads that krepp is able to map are not placed anywhere below the root (Fig. 4c): The portion of reads mapped by krepp that are successfully placed on the tree is significantly less for EMP compared to HMP.

### krepp improves taxonomic assignment on CAMI-II

We next tested if krepp distances can be used for taxonomic binning instead of phylogenetic placement, adopting it in two ways. The most straightforward approach is to assign each query the same taxonomic label as the reference with the minimum distance (krepp-closest hereafter). Since this approach ignores relations between taxonomic groups (represented as a multifurcating tree), we also directly applied the placement algorithm from krepp to the taxonomic tree. Marking “species” the lowest level of the taxonomic tree, we have a multi-furcating backbone tree. To compute krepp distances to each species, represented by one or more genomes, we compute the average Hamming distance histogram (**v**), mismatch count (u), and subsampling rate (ρ) for all genomes corresponding to it. With these average statistics at hand, and recalling that the krepp algorithm does not need a binary tree, we place each query independently on the taxonomic tree, noting that each branch corresponds to a taxonomic label at some rank. To benchmark krepp’s binning performance, we used the CAMI-II benchmarks [47], focusing on the taxonomic binning of pooled contigs obtained from the gold-standard assembly on their marine and strain-madness datasets. We used our largest index built from 123,853 genomes in the RefSeq snapshot provided for the challenge.

Both versions of krepp performed extremely well according to multiple metrics (e.g., accuracy, completeness, and purity) in both datasets across all taxonomic ranks (Fig. 5). Overall, krepp with the placement algorithm (krepp hereafter) exhibits the best binning performance, except at the species level for the marine dataset, where it is slightly behind the previously leading method Kraken2 [49]. No method other than Kraken2 is remotely competitive with krepp. Averaged across all ranks, krepp has 2.5% and 1.3% higher accuracy than Kraken2 in marine and strain-madness datasets, respectively. krepp-closest performs similarly to krepp in terms of accuracy, with differences only at the species level. For the strain-madness dataset, where many closely related strains can be present, krepp does better than krepp-closest, while the opposite is true on the marine dataset. Similar to the trends observed in our previous experiments, krepp and krepp-closest have the highest percentage of assigned sequences, with a considerable increase over the second-best method, Kraken2 (Fig. 5b). Beyond accuracy, krepp achieves noticeably higher completeness and purity than Kraken2 consistently in the strain-madness dataset (Fig. 5c). Although Kraken2 has marginally higher purity in the marine dataset, krepp compensates by having a commensurate increase in completeness, except at the species level. As expected, krepp creates more pure binning than the more aggressive krepp-closest in almost all conditions.

**Figure 5.**
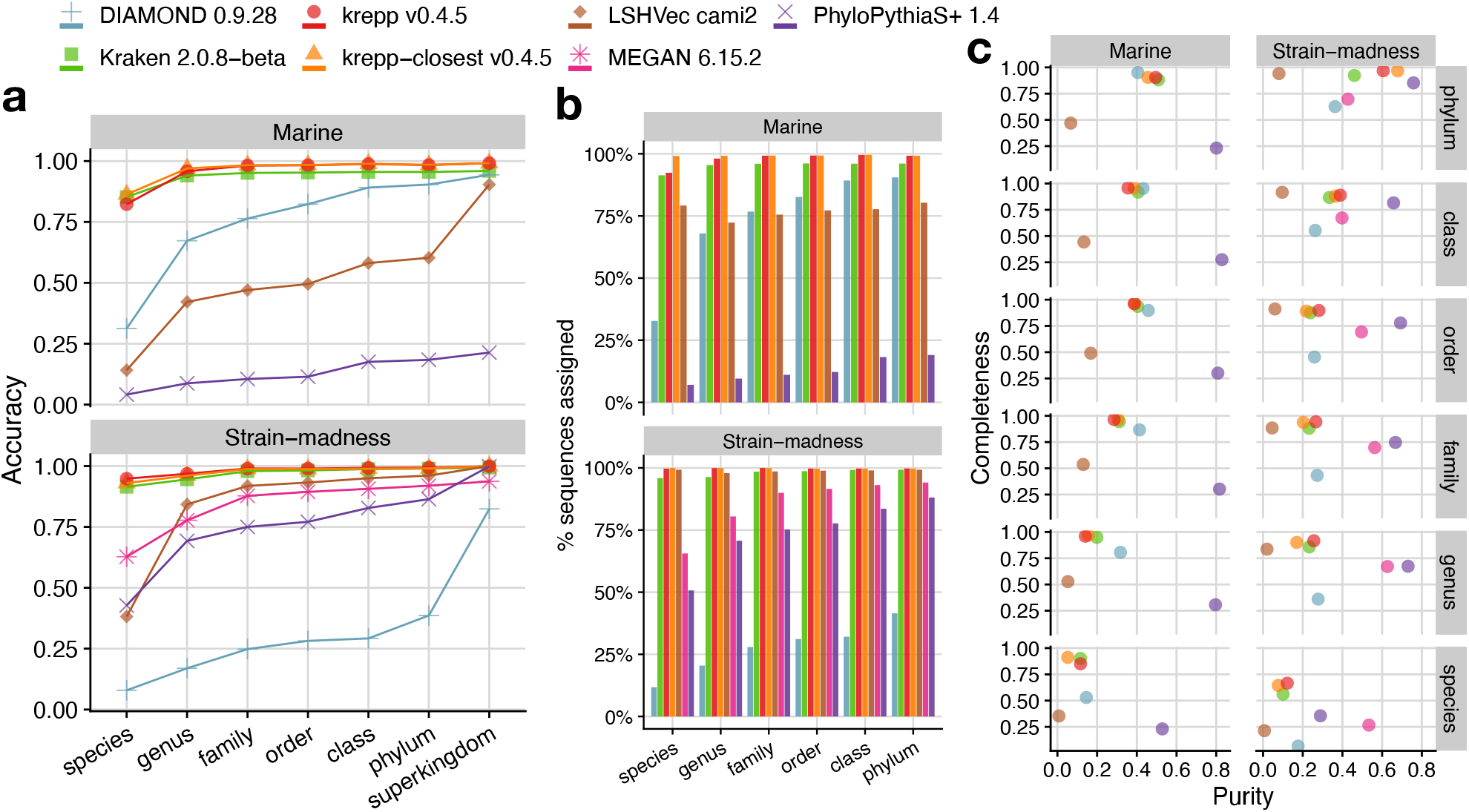
Accuracy (**a**), percentage of assigned sequences (**b**), and completeness versus purity (**c**) across taxonomic ranks in taxonomic binning of contigs of the gold standard assembly of CAMI-II. All metrics computed using AMBER [48].

## Discussion

We introduced the first scalable genome-wide phylogenetic read placement method and, in doing so, also designed a method to accurately estimate the distance of a read to a diverse set of genomes. Our method, krepp, provides a practical tool for performing new analyses of modern microbiome data with manageable computational requirements, enabling us to go beyond marker genes while maintaining high accuracy for phylogenetic placement. Integrating krepp with widely used pipelines that currently use alignment to map reads to operational genomic units (e.g., Woltka [21]) demonstrated superior performance in differentiation of real metagenomic samples from various environments, including host-associated (human or non-human) and free-living microbial communities.

As the number of references increases, the running time scalability of krepp (Fig. 2g) makes it more attractive than alignment, allowing utilization of larger reference datasets (e.g., a RefSeq snapshot with >120,000 genomes) to increase resolution. Scalability is achieved through LSH-based k-mer hashing, which limits the number of k-mers comparisons. To scale to even larger references (e.g., 200,000 genomes in GG2 [50]), running time will not be an issue, but memory could become a limiting factor. For such cases, krepp already supports a more aggressive filtering of k-mers using the unbiased methods FracMinHash [51, 52]; the extent to which k-mers can be subsampled while keeping high accuracy should be studied in the future. Future work can also incorporate tree-based methods (e.g., KRANK [27]) for k-mer subsampling.

Beyond k-mer selection and matching, alternative placement algorithms can also be explored. Our placement algorithm is currently branch length agnostic, enabling it to be applied with any binary or multifurcating tree, regardless of how it is computed. Moreover, its placements are interpretable (sister to the largest clade with statistically indistinguishable distances). However, it lacks formal guarantees for trees with high degrees of deviation from ultrametricity. When the backbone tree has branch lengths in interpretable units, we can take advantage of such lengths in the placement algorithm, a topic that we will explore in the future. Such methods could, in particular, address cases with very high levels of deviation from ultrametricity, perhaps using methods such as least-squares error placement [53, 54]. Conversely, our current algorithm needs a rooted tree, a restriction that should be ideally lifted since rooting trees is not always trivial. Finally, krepp can provide multiple placements per read, with associated probabilities. Future work should incorporate these measures of uncertainty in the downstream applications, including sample differentiation.

The framework designed here is general and can easily be adopted in other applications. The accurate read-genome distances can also be used for homology mapping [e.g., 55], taxonomic abundance profiling [e.g., 26, 42], contamination detection [e.g., 56, 57], contig-to-contig binning, computing rates of evolution across the genome to compute abnormally fast or slow regions (ultraconserved elements), and perhaps detecting horizontal transfer. Out of practical necessity, we used a species tree in our analyses, and this tree was sufficient for downstream applications such as sample comparison. We note that our placement errors have a long tail of high values, some of which correspond to low distances; a likely explanation is horizontal transfers from genomes that on the species tree are far from the query on the species tree, leading to gene tree discordance [58]. This observation can be used in the future to design methods that detect HGT or contamination across the genome.

## Methods

### Locality-sensitive hashing (LSH) index of *k*-mers

Given a query k-mer, we seek reference k-mers within some HD threshold, denoted by δ. Expanding on the CONSULT family of methods (CONSULT*) [25–27], we use LSH with some changes. We use the bit-sampling [59] to partition reference k-mers (default: k=29) into subsets. The LSH of a k-mer x, denoted by LSH(x), is computed by sampling h ≪ k (default: 14) random but fixed positions of x, providing [0, 2^2*h*^) buckets indexed by a 2h-bit integer. For each r ∈ ℛ, we use minimizers by choosing the k-mer whose encoding has the smallest MurmurHash3 [60] value in a local window (default 35). We save all surviving reference k-mers, denoted by ℳ, in the ascending order of their LSH(x), breaking ties lexicographically. The result is an array **A** of size |ℳ|. We build another ordered offset-index array of size 2^2*h*^, denoted by **I** to note the boundaries of LSH partitions; i.e., **I**[i] = **I**[i − 1] +|{x : LSH(x) = i, x ∈ ∈}| and **I**[− 1] = 0. Thus, unlike CONSULT* methods, which limit the size of LSH buckets by removing k-mers, krepp uses flexible size partitions and keeps all k-mers, which is helpful for distance calculations. To compute the exact HD for k-mer matches, **A** needs to store each k-mer x precisely. Naively, each k-mer requires 2k bits; however, the position of x in **A**, together with offset values in **I**, already gives the h positions used for LSH(x); we simply store the remaining 2(k − h) | |bits. Thus, **A** requires 2(*k* − *h*)| ℳ| bits. **I** is much smaller and needs log_2_(|ℳ|) per index (we write log(x) instead of ⌊log_2_(x − 1) ⌋ + 1 when clear by context) and 2^2*h*^ log(|ℳ|) bits in total. We adopt a left/right k-mer encoding [25] that enables computing HD with just four instructions (pop-count, XOR, OR, shift).

Given a query k-mer x, we only attempt to match it to k-mers with the same LSH value (i.e., **A**[**I**[LSH(x) −1]] to **A**[**I**[LSH(x)]]) by calculating the exact HD. The higher the h is, the smaller these slices tend to become, but with decreased sensitivity (i.e., more false negatives), especially for higher HD. However, since we explicitly calculate the HD, there are no false positives. Assuming independence of positions, two k-mers at HD = d have the same hash with probability 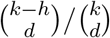 . This probability is sufficiently high for small enough d (e.g., d ≤ 4), then drops quickly and diminishes when d ≫ 4 for appropriate choices of h and k. For a query sequence of length L, the expected number of matches across all (L − k + 1) k-mers is sufficiently high for several realistic choices of k and h. The false negatives can be further reduced by using multiple arrays with different LSH functions (randomly sampled h positions), a feature that CONSULT* methods use, but we have not tested for krepp. Finally, note that k-mer matches at very high HD can be spurious (e.g., not orthologous) and will also have many false negatives as a query k-mer and the closest reference match are likely to have different LSH values. Thus, we choose a fixed parameter δ (default: 4) and only keep k-mer matches with HD ≤ δ.

### Coloring *k*-mers using a multitree based on the reference phylogeny

After finding reference k-mers similar to a query k-mer using the LSH index, we need to track *which* references include each matched k-mer x (denoted by ℛ (x) ⊆ ℛ). It is easy to save one ID for x, which is what many methods do (e.g., Kraken [9, 49] tracks the lowest common ancestor (LCA) of ℛ (x) while CONSULT-II stores a soft-LCA [26]). To calculate distances to all relevant references, we need to record all ℛ (x) IDs. Keeping pointers from x to each r ∈ ℛ (x) requires too much memory, necessitating an efficient data structure, which is the colored k-mer representation problem [28]. The high-level goal is to compactly represent all colors that are observed in a reference dataset and to retrieve them efficiently during query time for matched k-mers. A *color*, denoted by C, refers to a subset of references that share at least one k-mer (ℛ (x) for some x ∈ ℳ), and the goal is to compactly represent the set of all colors, denoted by 𝒞 = {ℛ (x) : x ∈ ℳ } . For simplicity, we assume all singleton colors ({r} for r ∈ ℛ) are in 𝒞 . Representing each non-singleton *C* ∈ 𝒞 as the union of other colors in defines a DAG G = (V, E) with colors as nodes and edges representing the partitioning. G can be stored in an array of size E + V (by saving the count and indexes of children of each node). The trivial bipartite DAG partitioning each non-singleton color into all its constituents will have |E| = ∑_*C*_∈|𝒸 | |C| − |ℛ| . However, we can do better by adding new nodes (i.e., “meta”-colors) for shared patterns across observed colors, which can potentially reduce the size (w.r.t. |E| + |V |). With meta-colors, we can further reap extra benefits from allowing exactly two children per node. Such a DAG can be stored in an array with 2|V | (instead of 3|V |) elements, each log(|V |) bits; the array index is the ID of the color, and for each index, we store the IDs of its children. For each x, we keep the index of ℛ (x) in an array **C** laid out identically to **A**; this adds log(|V |) bits to 2(k − h) bit k-mer encodings, which is manageable (e.g., |V | ≈ 2^22^ for our WoL-v2 reference |ℛ|= 15, 953). Additionally, we restrict the children of each node to be disjoint, making our DAG a binary multitree.

We argue that the phylogeny T provides a practical and efficient way to build the multitree. Consider a simple evolutionary model. An ancestral genome evolves down T accumulating random substitutions under an infinite k-mer assumption (similar to infinite sites) where the probability of any k-mer mutating twice is zero. Under this model, each ℛ (x) becomes a perfect character, meaning that it maps exactly to a single clade of T . As a result, all the colors in 𝒞 will be a node in T ; it is easy to see that the optimal solution w.r.t |E| is simply T after contracting internal nodes without a color associated (i.e., removing branches where no k-mer mutated). While this model is reasonable for relatively short time scales, genome evolution across phylogenetic scales is far more complex and does not produce perfect characters. To handle this complexity, we build a multitree instead of a tree.

We start with T as the multitree, representing each internal node as a color or meta-color. For simplicity, we describe our algorithm for bifurcating trees, noting that extending to multifurcating trees is straightforward. We partition each ℛ (x) into the set of clades C_1_, …, C_*n*_ ⊆ ℛ (x) of T such that no two clades C_*i*_ and C_*j*_ are sister to each other (i.e., all clades are maximal); note that each C_*i*_ can be a singleton. Note that n = 1 for a perfect character and n = |ℛ (x) | in the worst case. Each *C*_*i*_ is already a node of the multitree and thus can be represented with no additional color. For any set of n maximal clades, there can exist at most n − 1 nodes of T that have at least one maximal clade under at least two of their children. For each such node corresponding to ℛ (x), we define a (potentially new) color as the union of the colors of its children. With this procedure, we obtain ℛ (x) at the LCA of all clades C_1_, …, C_*n*_. We implement this idea using Algorithm 1, which builds **A** and the multitree jointly, moving up the tree T in a post-order traversal (implemented with nested task-parallelism). We find all k-mers shared between children, and for each such k-mer, we create a new color from its existing colors (line 19). The only difficulty is checking if the union of two colors already exists (Parent), which we implement using an Abelian group (details in the Supplementary Note). Adapting Algorithm 1 to a multifurcating node is done by building and adding indices of its children one by one in an arbitrary order (equivalent to resolving the node as a random ladder tree). For each child, we find its shared k-mers with all k-mers of previously added children and create new colors by repeating lines 12–21.

Our procedure adds n − 1 auxiliary colors on internal nodes, some of which are expected to be shared with other k-mers. This heuristic can be considered effective if a small proportion of the n − 1 added colors remain unobserved after we process all k-mers. We empirically observe this pattern; for 74% of non-singleton colors, none of their n − 1 added colors remain unobserved, and for 90% of colors, only ^1^/_3_ or less are unobserved (Fig. S1b).

### Maximum likelihood estimation of the distance

For a query read q, we start with all reference genomes with at least one k-mer match with HD capped at δ, filter out highly distant genomes when low distant ones also exist (see the Supplementary Note), and let the remaining reference genomes be ℛ*′*. We compute the distance of q to each r ∈ ℛ^*′*^. We make two simplifying assumptions: i) The k-mer match with the lowest HD is the orthologous one, and therefore, if multiple k-mers in r match a query k-mer, we take the lowest HD and discard others. ii) There is no dependency between adjacent k-mers, and thus, we ignore positions of matched k-mers. Overlapping k-mers are clearly not independent, and Blanca et al. [36] have modeled the dependencies for exact matches. However, analyzing the dependency between inexact k-mer matches is far more challenging. Thus, we adopt this simplifying assumption, noting that the use of minimizers reduces the number of overlapping k-mer matches and reduces dependence. Due to the independence, the likelihood function for distance D ∈ [0, 1] becomes

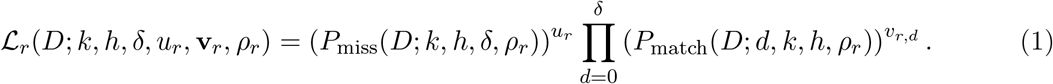

To compute P_match_, note that if we assume every match is due to homology, observing a match requires three independent events: i) The k-mer should be indexed despite using minimizers (or any form of random k-mer subsampling); we precompute this probability and call it ρ_*r*_ (see below). ii) Observing d mismatches when the underlying distance is D; this happens with probability

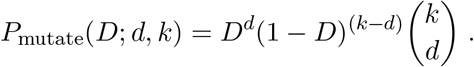

iii) The LSH search finds the k-mer match at HD = d with probability

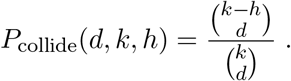

Therefore,

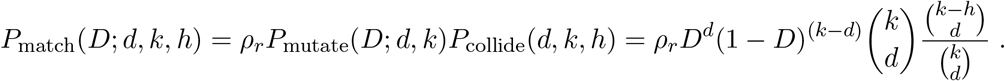

For ρ_*r*_, note we exclude k-mers only due to the use of minimizers. We precompute ρ_*r*_ during the indexing, setting it to the ratio of the number of minimizers to distinct k-mers for each r.

We can similarly compute the probability of missing a k-mer match, which can happen due to either of two disjoint events: i) The k-mer is not indexed, which happens with probability

1 − ρ_*r*_. ii) The k-mer is indexed but is not found by our method. An indexed k-mer can be missed either because it has HD > δ, and we automatically ignore such matches, or LSH does not find it (a false negative) even when HD ≤ δ. The probabilities of these two events are respectively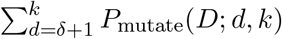 and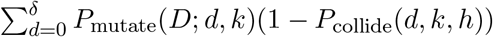 . Therefore, we end up with

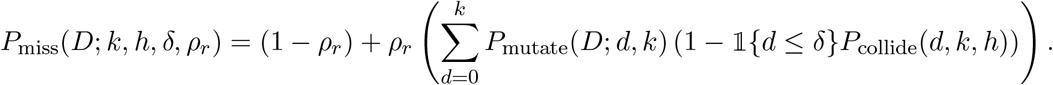

For each reference r, we maximize its likelihood function to get an estimate for D. Equivalently, we maximize the log-likelihood, which considerably simplifies Eq. (1) and P_match_ but does not help with the P_miss_ due to summation. After dropping constant terms (w.r.t D), we get:

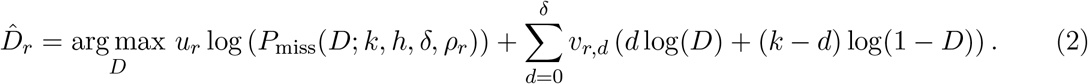

The log-likelihood function (Fig. 1D) is concave for D ∈ (0, 0.5) provided that k, h, and δ follow restrictions given in the Supplementary Note, including for the default values. We use the simple Brent’s method [61] for solving the optimization, which in preliminary analyses was faster than alternatives, such as L-BFGS-B [62]. Brent’s method uses quadratic interpolation to locate the global minimum by numerically approximating the derivative of single-variable convex functions.

#### Algorithm 1

Building the k-mer index (LSH index and multitree), given k-mers ℳ (r) for references r ∈ ℛ and the reference phylogeny T . ℳ_*i*_(r) is {x : x ∈ ℳ (r), LSH(x)=i}. We implement colors and C_1_ ∪ C_2_ (Parent) using integer encodings and an Abelian group hashing.

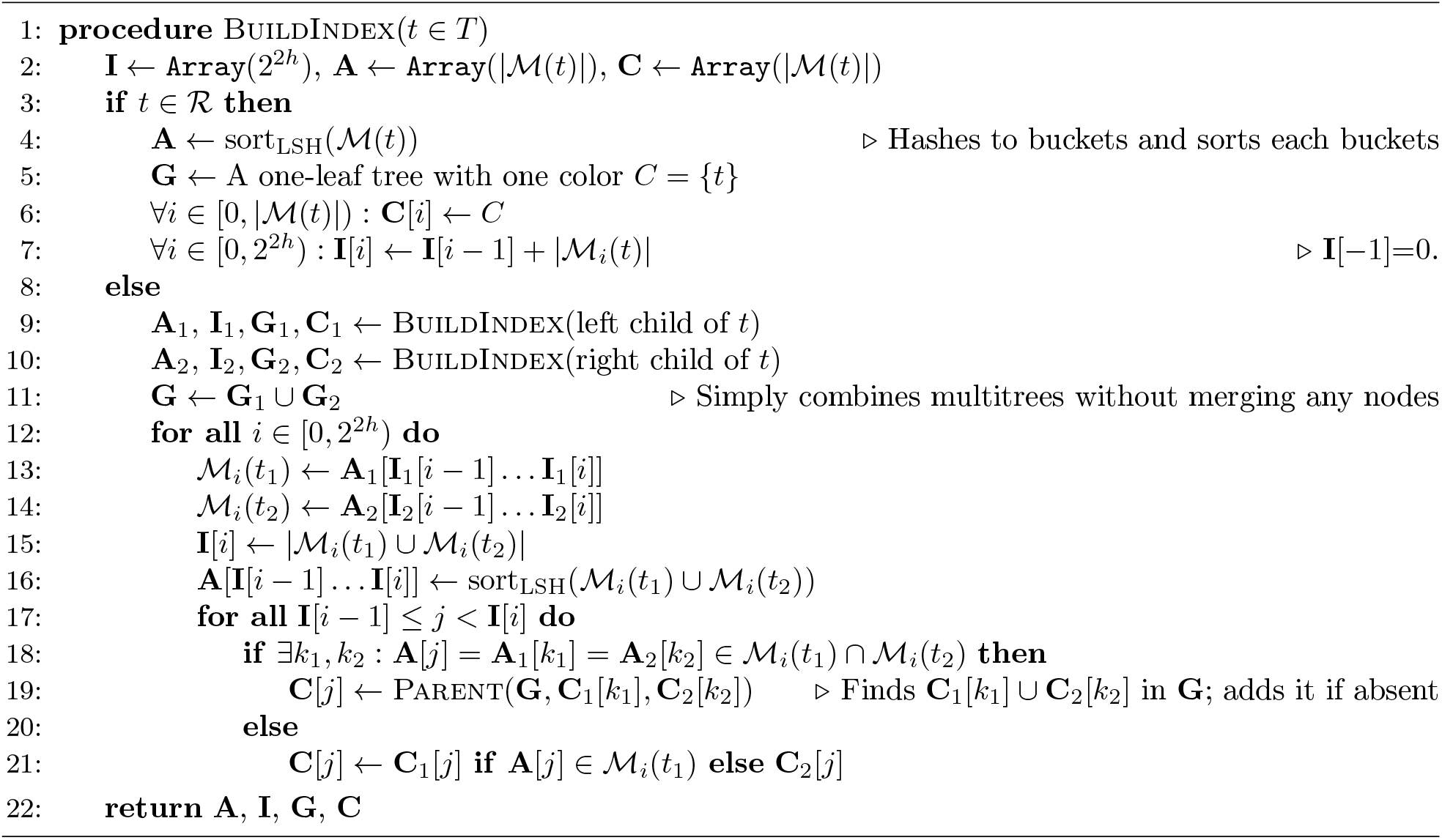

### Placement on phylogenies and taxonomies using likelihood-ratio test

When T is ultrametric, the distance from q to all leaves of its sister clade C is the same and is minimum across the tree (Fig. S7a). Therefore, given true distances, a valid approach is to place q as the sister to all references that have the minimum distance to it. This idea remains reasonable when the tree is not fully ultrametric but is sufficiently close to ultrametricity. Since distances calculated from reads have low resolution and high variance, we also need to consider that distances to many reference genomes may be statistically indistinguishable. Although one can consider using the least squares error minimization approach of APPLES [63] in this framework, we opt not to as it faces a subtle challenge (see the Supplementary Note) related to variable rates of evolution across the genome. We need methods that consider the high variability and noise of distances. We propose a principled likelihood-based test of statistical distinguishability for distances. With such a test, our goal becomes: Find the largest clade C of the reference tree T, such that the distance of q to references in C is statistically indistinguishable from the minimum distance of q to any node. We base our algorithm on this idea with a particular definition of clade distances (see below). Our formulation needs a root, which is often available and can otherwise be easily obtained [64].

#### Distance to a clade

On a bifurcating tree with bounded deviation from ultrametricity, q has low mean distances to both left and right children at the correct placement edge or any edge below it; on other nodes, it will have a high distance on one child and a lower distance on the other (Fig. 1E). To capture this intuition, we extend the notion of distance to a clade C parent to clades C_1_ and C_2_ by first defining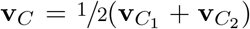. We also define 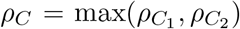. Similarly, for multifurcating trees (e.g., a taxonomic tree), we can define 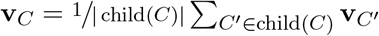 and ρ_*C*_ = max({ρ_*C*_*′* : C^*′*^ ∈ child(C) }) where child(C) denotes the children of C. With these definitions, Eqs. (1) and (2) are extended to clades and are used to compute the read-to-clade distances.

#### Statistical distinguishability

Small ML distances d_*∅*_ and d_*A*_ to two clades C_*∅*_ and C_*A*_ may differ due to random noise or small changes in ρ (a clade can be just a single reference). We can use likelihood to statistically test for this. Let C_*∅*_ be the one with the lower distance, and let l_*∅*_ and l_*A*_ be the log-likelihood (log of Eq. (1)) computed at d_*∅*_ and d_*A*_, using **v**_*A*_ as data. The likelihood ratio test (i.e., χ^2^ test with 1 degree of freedom on 2(l_*A*_ − l_*∅*_)) can be used to test if the higher likelihood clade is statistically distinguishable from the null (default α = 10% significance level).

#### Finding optimal placement

For every internal node of the tree, we compute the distance and use the indistinguishability test to ask if its distance is statistically tied with the minimum-distance clade. Among all clades where we fail to reject the null hypothesis, we choose the largest, breaking ties by choosing the clade with the smallest distance. We do not place reads if 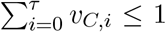 for the chosen clade C (default: τ = 2). Furthermore, if the chosen clade is the root, we characterize q as unplaced.

### Benchmarking

#### Genome evolution simulations

Starting from a genome X as the base, we added mutations to it at random using the Jukes-Cantor model with Gamma model of rate heterogeneity, with the added caveat that gene boundaries were respected (i.e., stop codons were not disturbed) and mutations fall only on the genes and not the ≈5% of the genome that constitutes intergenic regions. We then computed the genomic nucleotide distance (GND) from the mutated genome, X^*′*^, to X by counting the actual number of mutations added during the simulation, divided by the length (including intergenic regions). We mutated X at various nucleotide diversity levels (D ∈ {0%, 0.05%, 0.1%, 0.2%, …, 1%, 2%, …, 19%}) to obtain X_0_, …, X_30_. We then compared X with all X_*i*_ (i ∈ [0, 30]) genomes. For X, we used 29 bacterial single-cell assembled genomes from the GORG dataset (available at NCBI under bioproject ID GenBank: PRJEB33281) [37], selected to have high levels of completion. To create rate heterogeneity, we draw a relative rate multiplier r_*j*_ for each gene j of X from a Gamma distribution with mean 1 and variance ^1^/_*α*_ (here, α = 5), keeping rates the same across all levels. We assigned a fixed rate per gene to the base genome X, which is used for all X_*i*_, and then randomly selected n = D · L nucleotides (sampling without replacement) to mutate, selecting each position with a probability proportional to the rate r_*j*_ of the gene j it belongs to. We avoid adding or removing start and stop codons (to keep gene boundaries intact) and interrupting the reading frame. Overlapping areas between genes get mutated only if they do not change the start and stop codons. The rate multipliers of these overlapping areas are randomly selected among the genes that include them. Finally, using ART [65], we simulated 150bp error-free reads from 899 (29 × 31) mutated genomes at 1-fold of read coverage (setting `-f 1 -l 150 -s 10`), then subsampled to 4 million in total (each query contributing equally) using seqtk [66]. To compute the true Hamming distance, we used the coordinates of simulated reads.

### Distance and placement evaluation on WoL

For benchmarking of krepp distances with real genomes, we selected 500 query genomes across 36 phyla (267 genera) from RefSeq, with varying distances to the closest reference genome in WoL-v2 (Fig. S3a), quantified by the average nucleotide identity estimation (ANI) computed using Mash [24] (v2.3 with sketch size of s=100,000 and k=29). Then, we simulated 150bp short reads at high coverage with default Illumina error profiles using ART (setting -ss HS25 -l 150 -s 10), and subsampled to 33M reads in total. To select queries for phylogenetic placement, we clustered leaves of the WoL-v2 tree using TreeCluster [67] and set the length threshold parameter to the following values: 0.25, 0.1, 0.075, 0.05, 0.04, 0.03, 0.02, 0.01, 0.005, 0.001, 0.0005, and 0.0001. Each of these threshold values results in a clustering of leaves, where some remain unclustered as singletons. For all threshold values except 0.25, we selected all singletons that are not singletons with a larger threshold value. Among these, we sampled 10 leaves at random for each level, resulting in 110 query genomes we used for placement benchmarking with respect to the WoL-v2 dataset (Fig. 3a and Fig. 3b).

On these datasets, we compare krepp (v0.4.5) against distances obtained from alignment and placing on the closest tree leaf using bowtie2 (v2.4.1). For all bowtie2 analyses, to find all alignments with high sensitivity, we ran it with `--very-sensitive` and --all options.

We evaluated the accuracy of krepp’s placements in three different experiments, comparing krepp placements with placing as sister to the closest reference, comparing krepp and App-SpaM [11] on genome-wide placement of short reads from small clades, genome-wide placement (krepp) versus marker-based maximum-likelihood placement (EPA-ng [13]). In all experiments, we measure error by the number of edges between the output of the methods and the correct placement before pruning. We simulate genome-wide 150bp short reads as done in the previous analyses with default Illumina error profiles and attempt to place every read on the backbone after pruning the query genome(s). For 16S comparison (Fig. 3d), we selected 100 queries from WoL-v1 with varying novelty levels based on the closest neighbor in the backbone tree WoL-v1 (Fig. S3b). Then, we simulated error-prone short reads using the same ART parameters and subsampled to 1M reads for each of these two query sets. Out of 10,575 genomes in WoL-v1, 7813 genomes included a 16S gene. For EPA-ng, we placed 16S reads after removing all genomes without a 16S gene from the reference tree, whereas genome-wide read placement with krepp is performed without excluding such genomes.

#### Real microbiome analysis

Human microbiome samples analyzed consist of the following seven body sites: stool (n = 78), tongue dorsum (n = 42), supragingival plaque (n = 33), buccal mucosa (n = 28), retroauricular crease (n = 13), posterior fornix (n = 10), and anterior nares (n = 6). We used 1M 100bp paired-end whole-genome sequencing reads subsampled in Zhu et al. [21]. krepp processes paired-end reads separately and estimates distances independently. We built count-based BIOM tables [68] (Biological Observation Matrix) using a separate module available as part of Woltka (`woltka classify`) for krepp (see Supplementary Note). For Woltka OGUs and Bracken’s species-level profiles, we obtained UniFrac and Bray-Curtis distance matrices from Zhu et al. [21] (which were also calculated using WoL-v2 reference dataset) and used those for comparison. We used the BIOM tables, rarefied to 100,000 reads per sample after filtering relative abundances below 0.01% (removing two samples), to compute pairwise distances using weighted UniFrac (Woltka and krepp) or Bray-Curtis for taxonomy-based methods.

To test improvements when the dataset size increases, we built two new indices based on the ultra-large phylogenetic tree consisting of 199,330 reference genomes inferred using uDance by augmenting the WoL-v2 tree [3]. Among these genomes, we only retained the ones that also appear in the RefSeq snapshot (as of 2019/01/08) provided by CAMI-II benchmarking [47], resulting in 50,752 genomes, and pruned the original uDance backbone tree down to these genomes. Since Balaban et al. [3] removed near-duplicate genomes from the initially curated set of 656,574 microbial genomes, we further expanded the pruned uDance tree by adding near-duplicates from the intersection of this set and the RefSeq snapshot of CAMI-II. Making near-duplicated genomes sisters (with 0 branch lengths) resulted in the tree consisting of 123,853 references.

Earth microbiome samples consist of 746 microbial communities categorized into four levels (as shown in Fig. 4d). Raw shotgun sequence data for these samples is available through Qiita at qiita. ucsd.edu (study: 13114). The total number of reads was 14,720,788,401 (with a mean of 1,947,709 and a standard deviation of 5,446,907) across all samples. We placed all reads in these samples and estimated distances to find OGUs using krepp, then rarefied the BIOM tables obtained using `woltka classify` to 6,550 reads per sample (following Shaffer et al. [46]), resulting in 686 samples with sufficiently high sampling depth. Note that only 612 samples survived rarefication at the same sampling depth when the alignment was used, demonstrating the impact of the higher mapping rate of krepp. BIOM tables from outputs of all tools were constructed via woltka classify command and filtering was done via `woltka filter`, setting `--min-percent` 0.01 [21]. For PERMONOVA test and UniFrac computation, we used QIIME2 [69] and its diversity plugin [43], setting sampling depth to 100,000 for human microbiome and 6,550 for Earth’s microbiome, and the number of permutations to 1,000. For Earth’s microbiome, we report pseudo-F statistics computed in [46] using bowtie2, Woltka, and the same QIIME2 workflow.

### CAMI-II taxonomic binning of contigs

Reference genomes (a RefSeq snapshot as of January 8, 2019) and the taxonomy for CAMI-II experiments are available at cami-challenge.org/reference-databases/, contigs from the gold standard assembly can be found at frl.publisso.de/data/frl:6425521/marine/ and frl.publisso.de/data/frl:6425521/strain/, for marine and strain-madness datasets, 2.59 and 1.45 gigabases, respectively [70]. For all tools except krepp (DIAMOND [71], Kraken2 [49], LSHVec [72], MEGAN [73], and PhyloPythiaS+ [74]), we used CAMI-II results submitted to the challenge, available at (github.com/CAMI-challenge/second_challenge_evaluation). For krepp, we used the same reference set of 123,853 references used for HMP, noting that this set was already filtered not to include any genome not present in the CAMI-II reference set. We used AMBER (github.com/CAMI-challenge/AMBER) [48] to compute evaluation metrics (accuracy, percentage of assigned sequences, completeness and purity), exactly as also done by Meyer et al. [47] for other methods.

## Data availability

The list of krepp indexes used across our experiments can be found at ter-trees.ucsd.edu/data/krepp/. The Web of Life database can be accessed at biocore.github.io/wol/ (WoL-v1) and ftp.microbio.me/pub/wol2/ (WoL-v2). We made simulated genomes, all simulated reads, and auxiliary data, together with distance estimates, placement results, and BIOM tables, available on Dryad [75].

## Code availability

krepp, implemented in C++17 and optimized using OpenMP [76], is freely available at github.com/bo1929/krepp [77], together with a user manual and a tutorial. We used the Boost Math library [78] for numerical optimization, and the parallel-hashmap library [79] for hash maps. Key scripts for reproducibility (experiments, evaluation metrics, and figures) are also provided at github.com/bo1929/shared.krepp and on Dryad [75].

## Acknowledgement

This work was supported by the National Institutes of Health (1R35GM142725) grant and a Minderoo Foundation research grant to S.M. This work used Expanse at San Diego Supercomputing Center through allocation ASC150046 from the Advanced Cyberinfrastructure Coordination Ecosystem: Services & Support (ACCESS) program, which is supported by U.S. National Science Foundation grants #2138259, #2138286, #2138307, #2137603, and #2138296. We thank Qiyun Zhu for the support provided on the human microbiome analysis.

## Supplementary Figures

**Figure S1.**
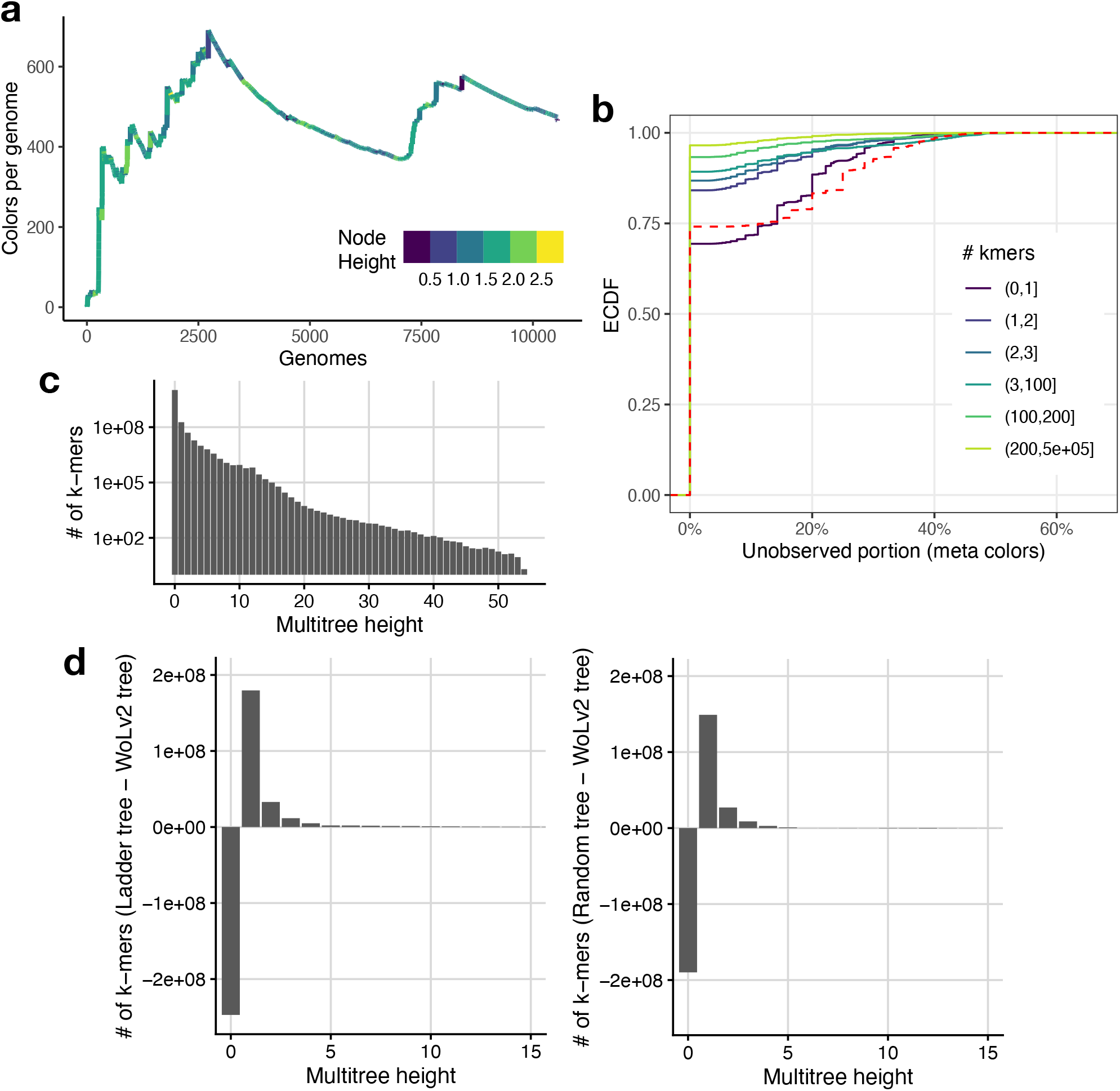
**a**, On WoL-v1 data set (with 10,575 genomes), the number of nodes per genome as postorder progresses (*x*-axis). **b**, For every color in the WoL-v2 reference dataset with 15,953 genomes, we visit the colors below it on the multitree; we then count what percentage of these colors are meta-colors (those not associated with any *k*-mer). We show the empirical cumulative distribution of the portion of colors under each color are unobserved. We divide this based on how many *k*-mers are labelled by each color, showing the undivided data in the dotted red line. **c**, The histogram of the number of *k*-mers with a certain height in WoL-v2 dataset. Most *k*-mers belong to colors with height 0, 1, or 2. **d**, Comparing the number of *k*-mers with colors with certain multitree heights when different trees are used. The differences in the number of *k*-mers between: a random ladder tree and WoL-v2 tree (left), a random dual-birth model tree (*λ*_*A*_ = 10, *λ*_*B*_ = 1) [80] and WoL-v2 tree (right) are shown.

**Figure S2.**
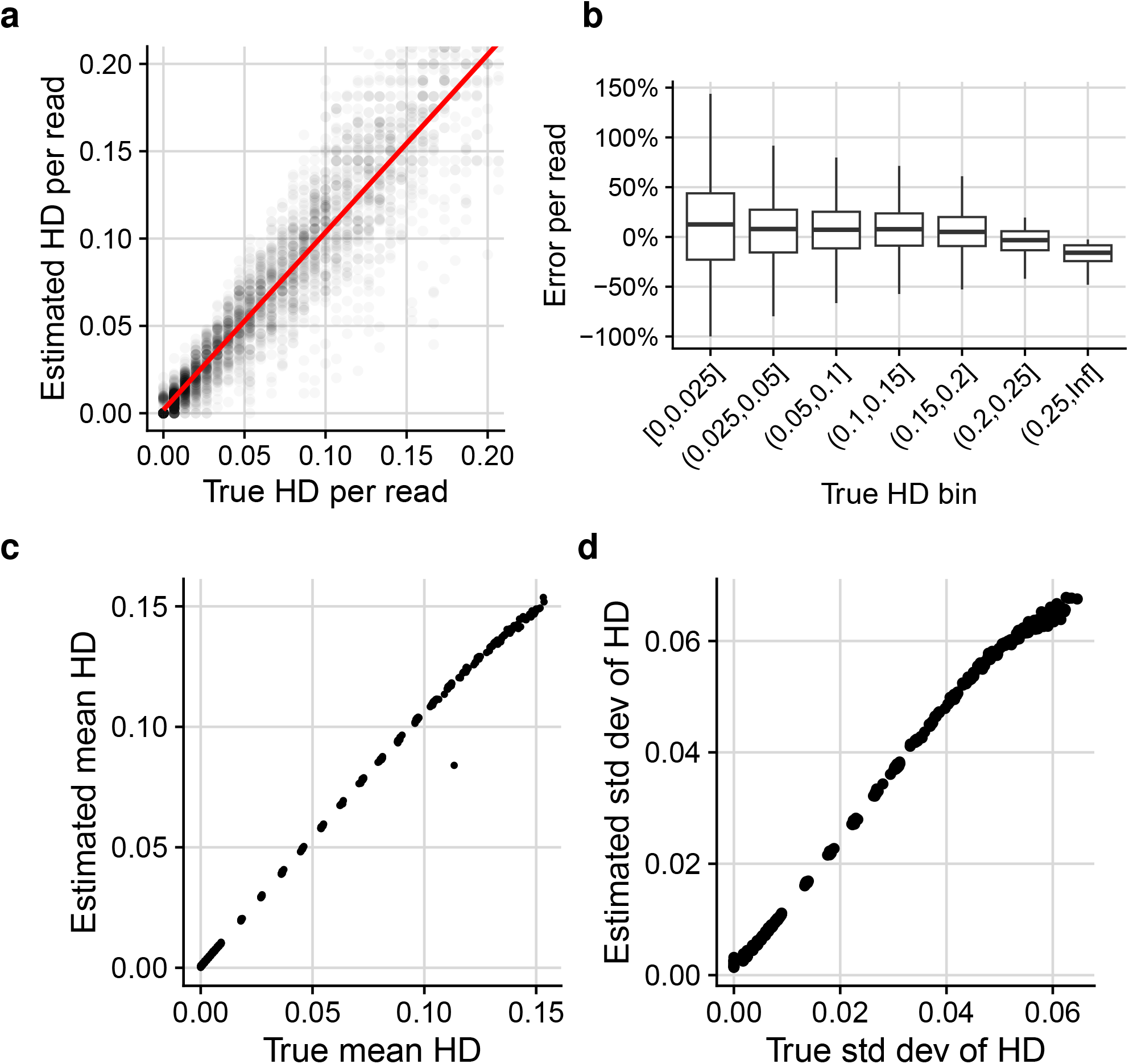
**a**, Comparing true Hamming distance (normalized by the read length) and the estimated HD for individual 150bp short reads. Each data point is a read and we plot 100,000 randomly subsampled reads in total. The red line is fit using a linear model. **b**, Read-level percentage error distributions for each true HD bin, demonstrating small overestimation bias, especially in low Hamming distances. **c, d**, Each data point is a mutated genome, and the mean and standard deviation values are computed across all reads that krepp is able to map to the corresponding base genome.

**Figure S3.**
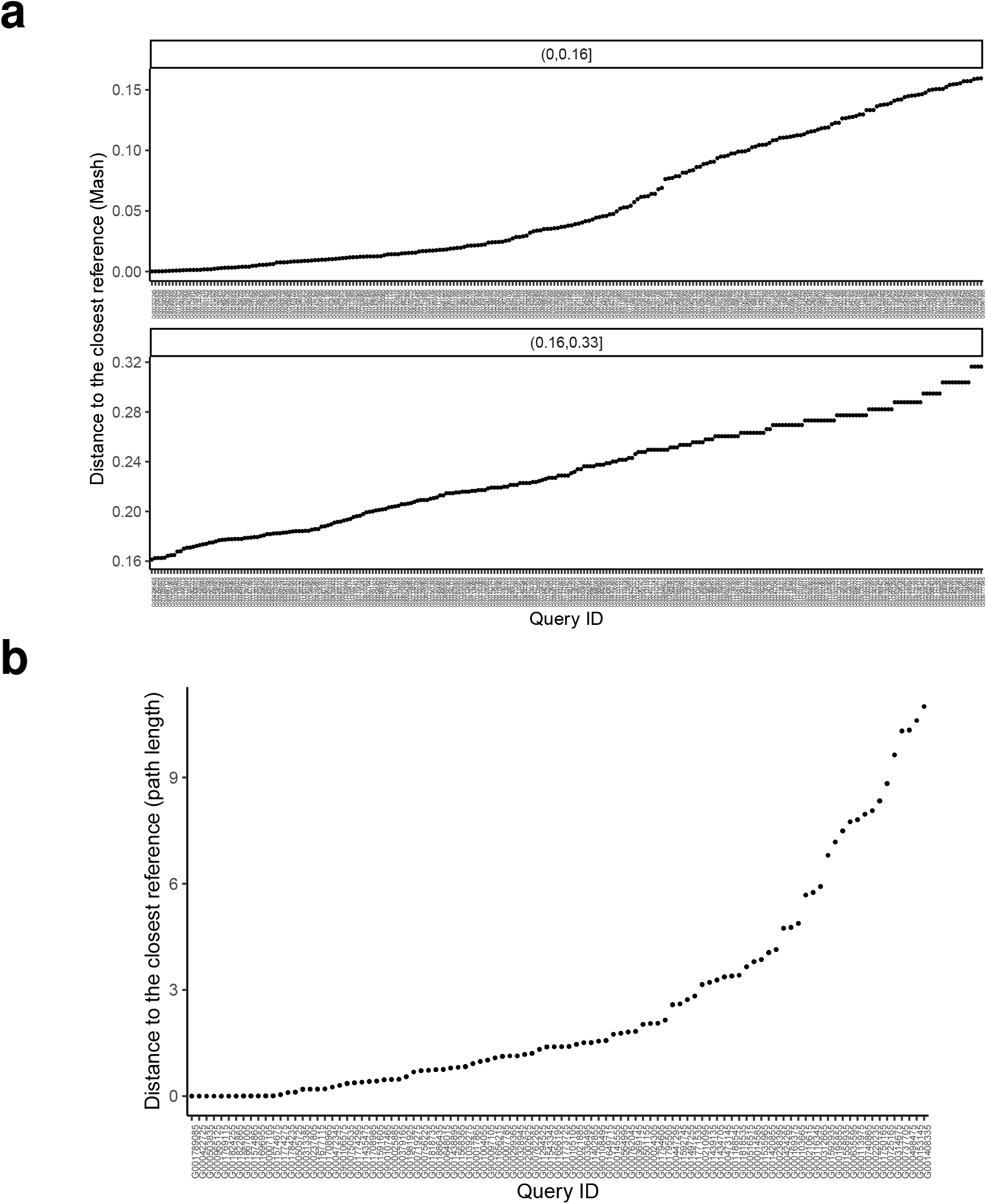
**a**, Distances of 110 queries analyzed for distance benchmarking, measured by 1-ANI to the closest reference in WoL-v2. **b**, Path lengths to the closest reference for 100 queries selected from WoL-v1 tree.

**Figure S4.**
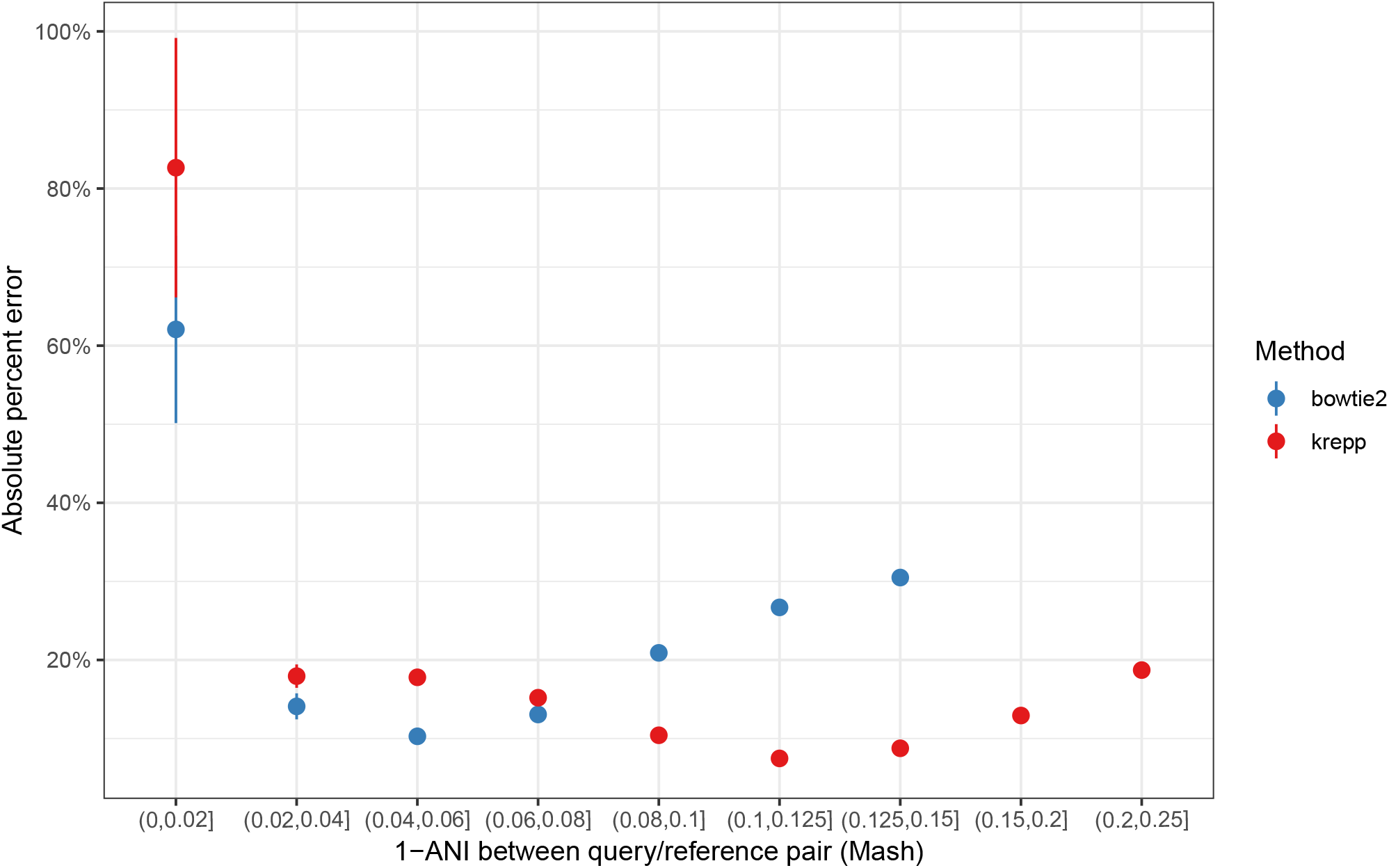
Change in the relative error 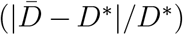 of krepp and bowtie2 across genome-wide distance bins (measured by Mash) for query/reference pairs with at least 20% reads mapped by each method.

**Figure S5.**
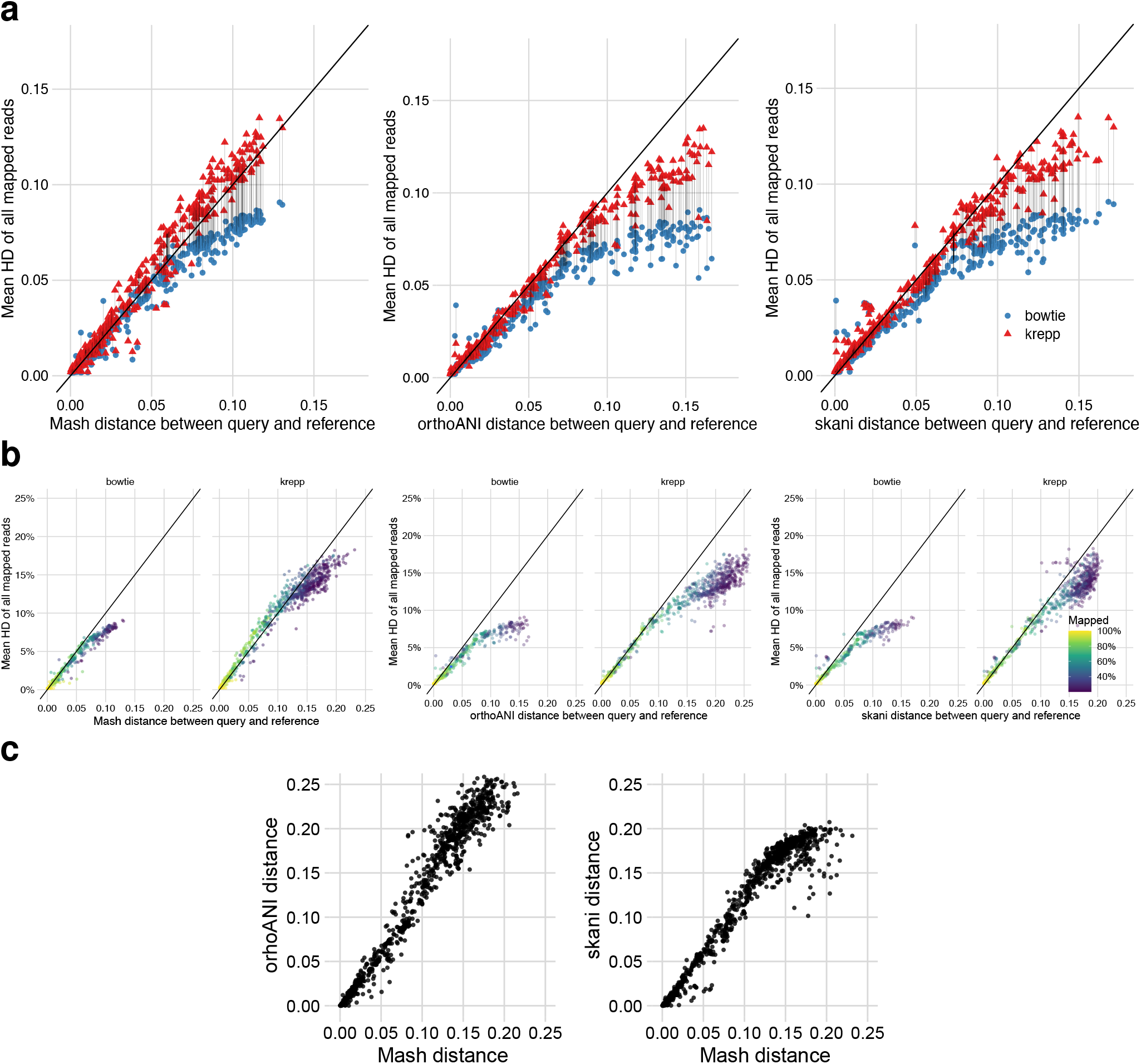
**a**, Mean distance across reads 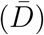 versus orthoANI [39] and skani [38] genome-wide distance estimates *D*^*∗*^ for query/ref genome pairs with ≥20% reads mapped by both methods. **b**, 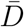 versus orthoANI and skani distance estimates *D*^*∗*^ from each query to all references with at least 20% reads mapped (colors). **c**, Comparing genome-wide ANI estimates of Mash with orthoANI and skani. We set `--min-af` parameter of skani to 0 to output distances regardless of the aligned fraction value. All other parameters of orthoANI and skani are set to defaults.

**Figure S6.**
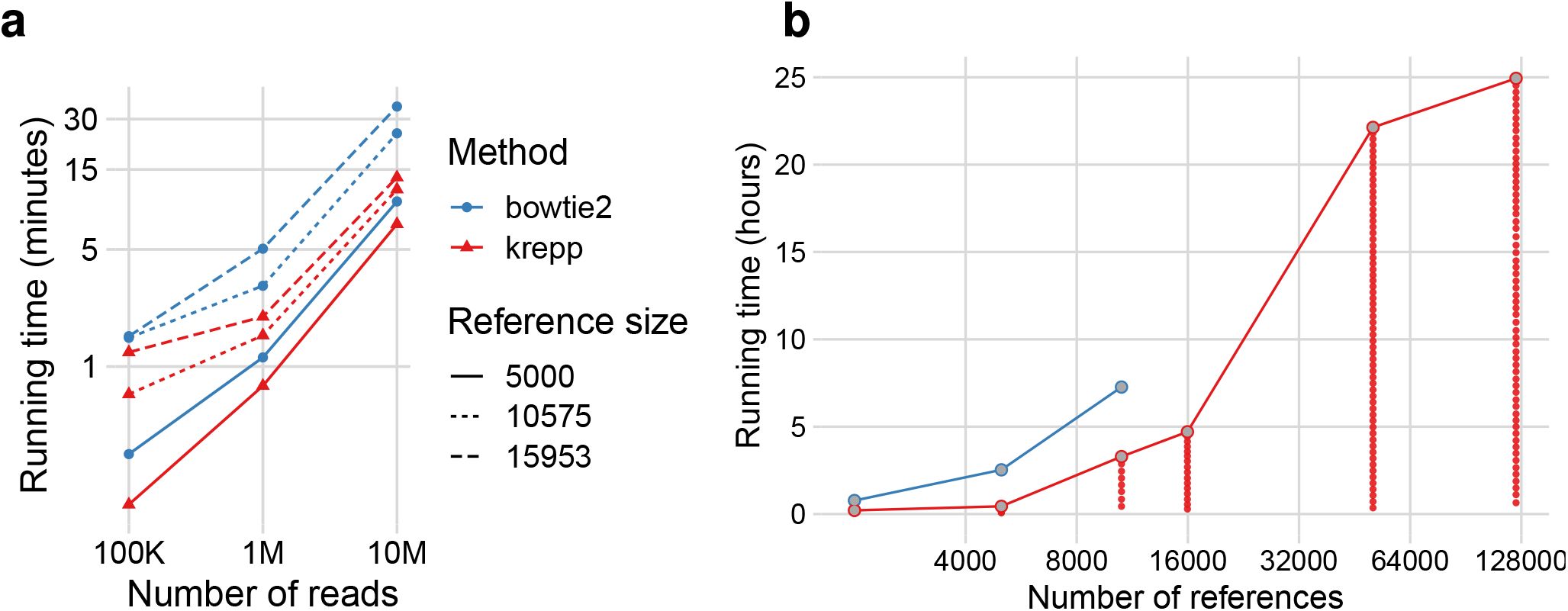
Scaling. **a**, Querying time versus the number of reads. Both methods scale similarly. **b** Indexing time versus the number of references. krepp builds the index in batches (a set of consecutive rows of the LSH index); we show the sum (line), but batches (dots) are run separately in parallel. WoL-v2 reference (15,953 genomes) with bowtie2 had to be built on a more powerful machine, taking 5.5 hours, and cannot be compared.

**Figure S7.**
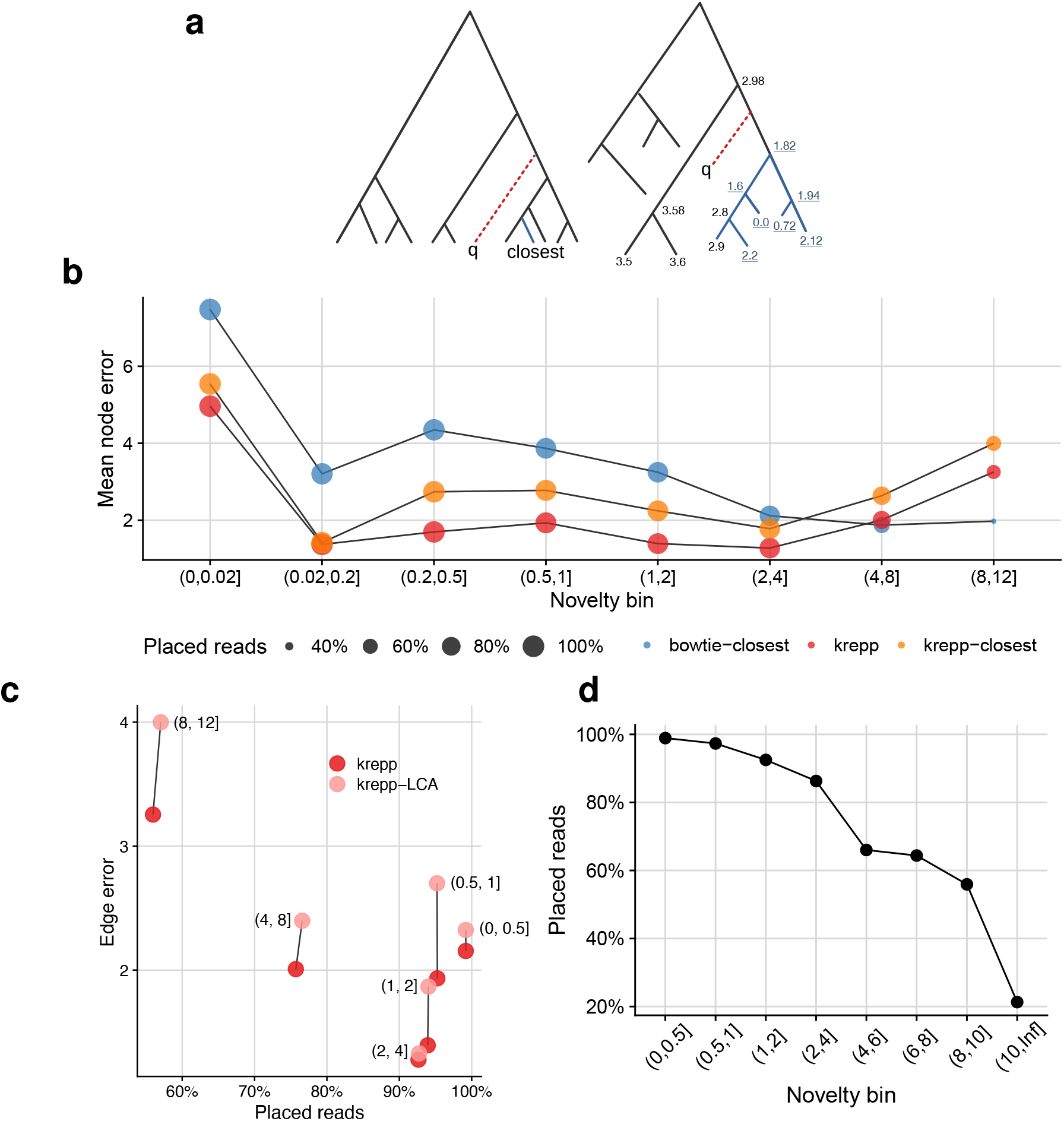
**a**, On an ultrametric tree, the query *q* has the same minimum distances to its sister clade; thus, the choice of the closest leaf is arbitrary unless the sister clade is a singleton. When the backbone tree is sufficiently close to ultrametricity, distances to the sister clade might be similar and statistically indistinguishable; placing the query as sister to the largest clade of similarly small distances can find the correct placement. Each node is labeled with its *χ*^2^ value according to our likelihood ratio test; all values below 10% significance are indistinguishable (underlined). **b**, Placement error (*y*-axis) for 110 query genomesm, binned based on novelty measured as the path length to the closest leaf on the WoL-v2 tree (*x*-axis). **c**, Using krepp distances and statistical test of distinguishability, we can find the set of leaves that are all tied with the closest distance to the query; the read can then be placed as sister to the lowest common ancestor (LCA) of these leaves. This approach increases the error compared to our default algorithm. **d**, Percentage of reads placed for queries selected from WoL-v1 across varying novelty levels. We observed reduced placement rate compared to using more densely-sampled WoL-v2 as the reference.

**Figure S8.**
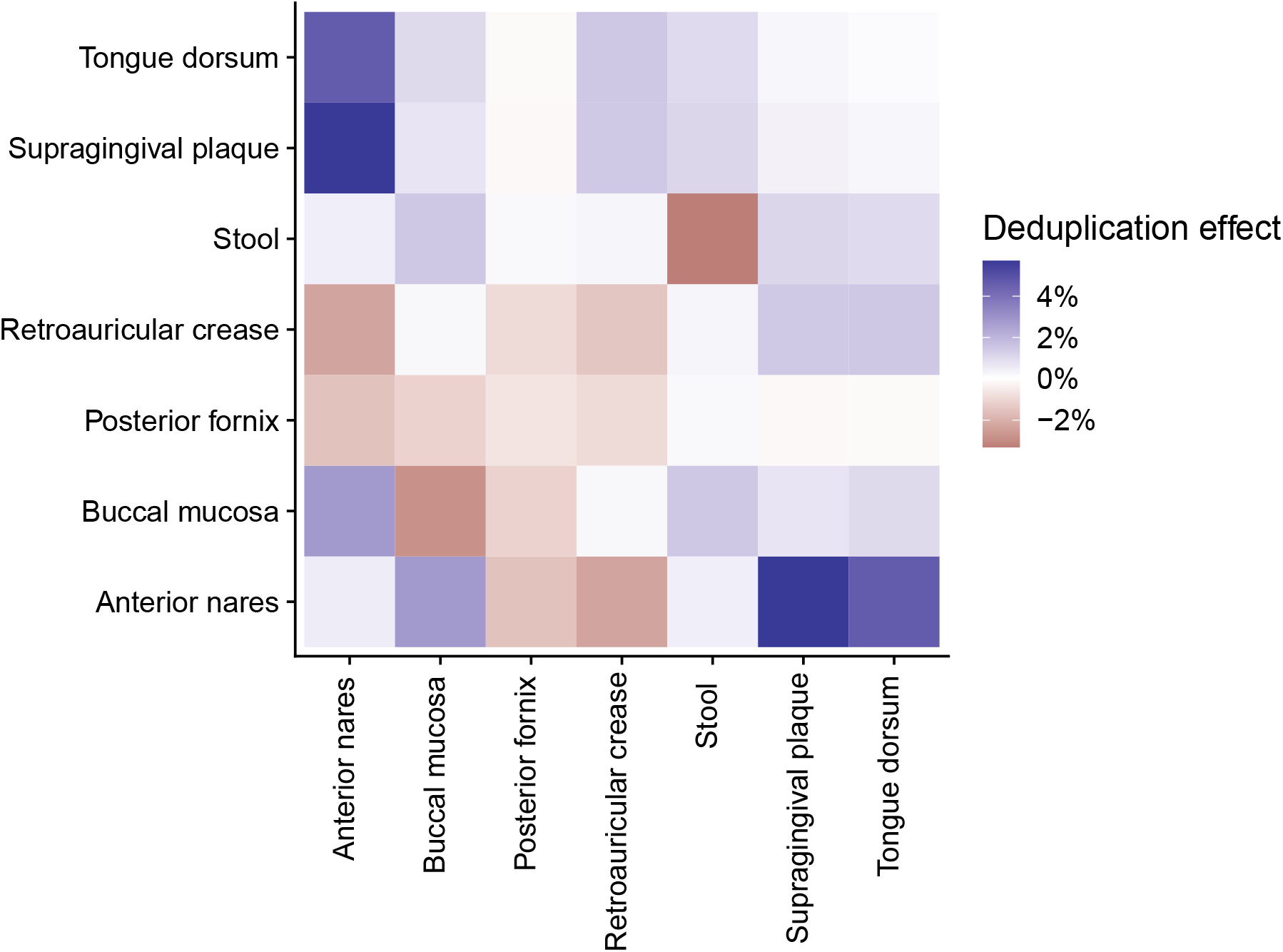
Effect of deduplication on weighted UniFrac distances across samples from all body sites. We show the average change in distances when krepp placements are computed with respect to the deduplicated RefSeq index, instead of the one with near-duplicates.

**Figure S9.**
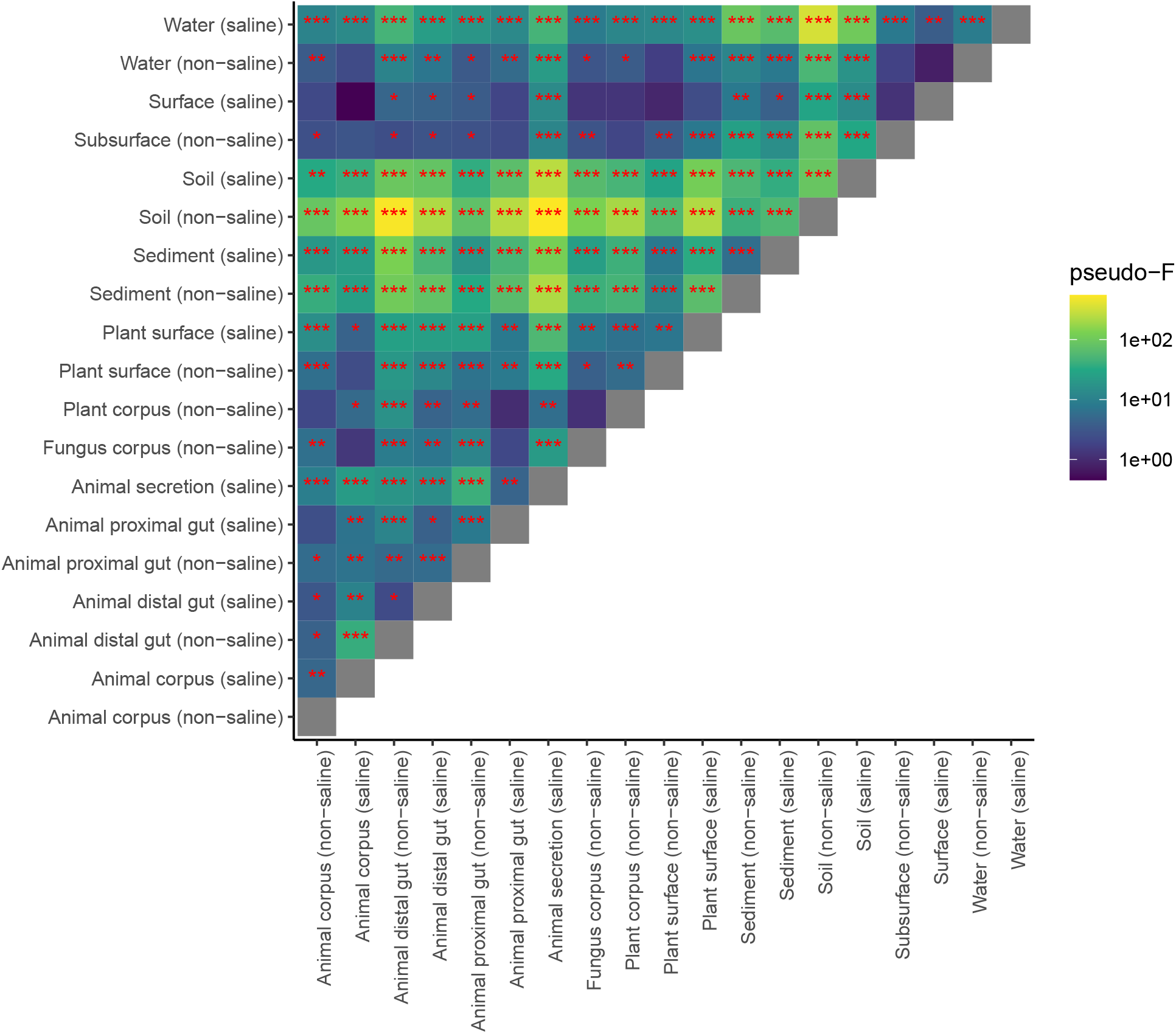
Pseudo-*F* and significance of separation based on pairwise PERMANOVA test across different environments at EMPO 4. The number of stars for each environment pair corresponds to different levels of *p*-value (shown as ∗: ≤ 5%, ∗∗: ≤ 1%, ∗∗∗: ≤ 0.1%).

## Supplementary Tables

**Table S1.** Notations used throughout the paper.

| Symbol | Definition | Symbol | Definition |
| --- | --- | --- | --- |
| $\mathcal{R}$ | Set of all references | $\mathcal{R}(x) \subseteq \mathcal{R}$ | Set of genomes that include a $k$-mer $x$ |
| $T$ | Reference tree with $\mathcal{R}$ as leafset | $\mathcal{M}$ | All $k$-mers in $\mathcal{R}$ after minimization |
| $\delta = 4$ | Maximum HD for $k$-mers | $\mathcal{C} = \{\mathcal{R}(x) : x \in \mathcal{M}\}$ | Set of all observed colors |
| $h$ | Positions sampled for LSH | $\mathbf{A}, \mathbf{C}$ | LSH sorted arrays of $k$-mers and color IDs |
| $\rho_r$ | Subsampling rate for reference $r$ | $\mathbf{I}$ | Offset-index array for bucket boundaries |

**Table S2.**
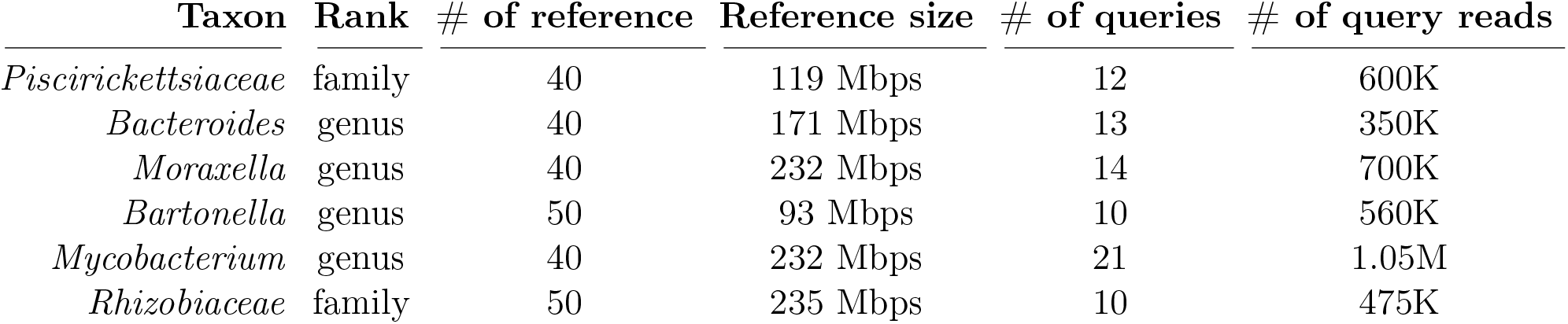
Details of small reference sets consisting of a single taxon and their corresponding query sets. Query genomes are simply the remaining genomes after excluding randomly selected genomes from WoL-v1.

| **Taxon** | **Rank** | **# of reference** | **Reference size** | **# of queries** | **# of query reads** |
| --- | --- | --- | --- | --- | --- |
| *Piscirickettsiaceae* | family | 40 | 119 Mbps | 12 | 600K |
| *Bacteroides* | genus | 40 | 171 Mbps | 13 | 350K |
| *Moraxella* | genus | 40 | 232 Mbps | 14 | 700K |
| *Bartonella* | genus | 50 | 93 Mbps | 10 | 560K |
| *Mycobacterium* | genus | 40 | 232 Mbps | 21 | 1.05M |
| *Rhizobiaceae* | family | 50 | 235 Mbps | 10 | 475K |

## Supplementary Note

### Efficient union of sibling indexes

#### Algorithm 2

Building the k-mer index (LSH index and multitree).

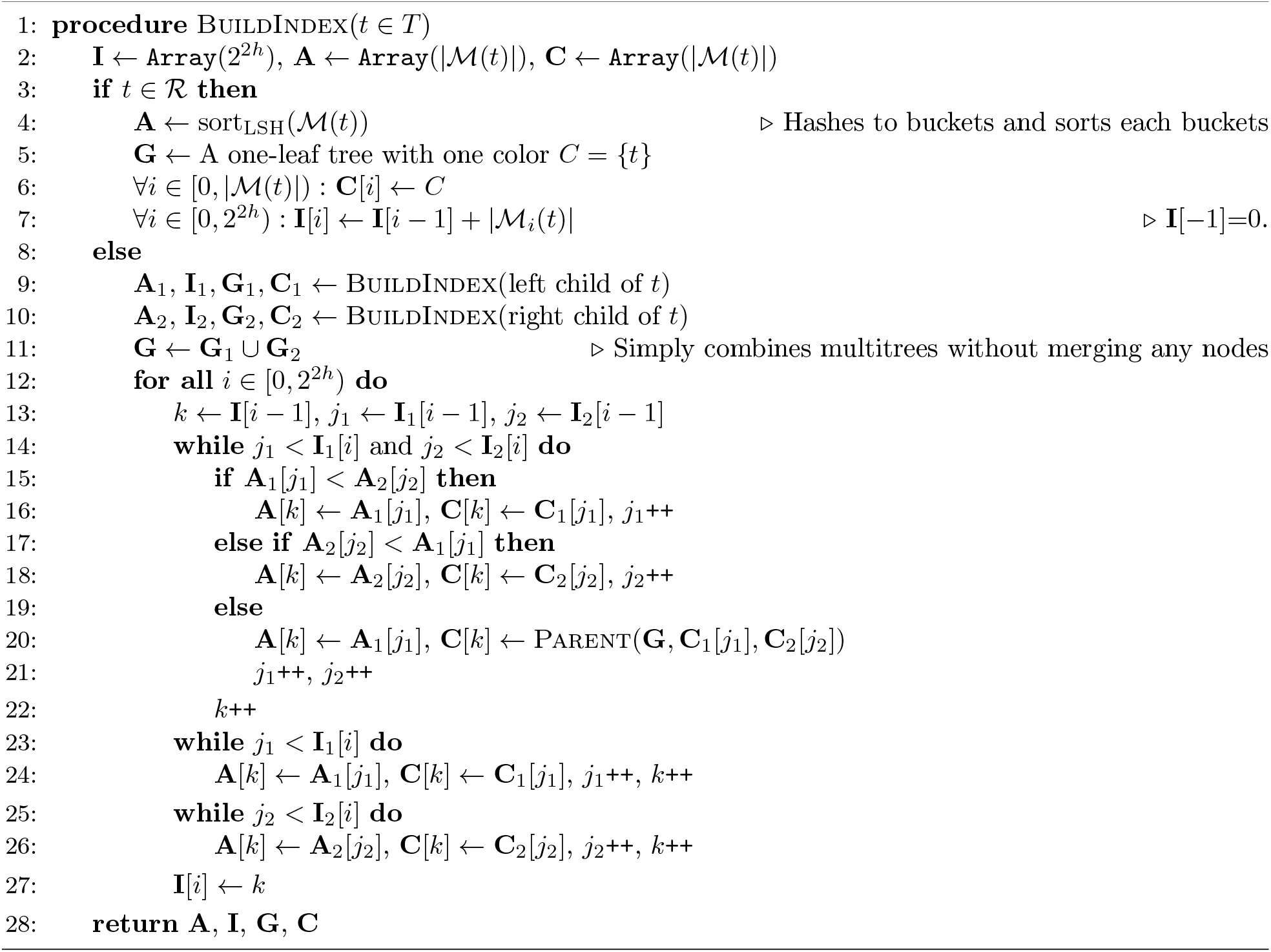

### Filtering high distance genomes and placements

We perform the following forms of filtering before computing distances.

- When a read q comes from a region that is shared across many references, the number of matches can be excessively high, especially for high δ (e.g., > 2). For the sake of running time, we ignore references with only high HD matches if low HD matches exist for other references. Specifically, let d_min_ be the minimum HD across all k-mers for all r ∈ R. We ignore a reference if the minimum HD across all its matches to q is 2(d_min_ + 1) or higher.
- Optionally, krepp can only report matching references with distances statistically indistinguishable from the ML distance of the closest reference, using the same likelihood ratio test as the placement. This option is disabled by default.

After computing distances and choosing a candidate clade C (the largest one that we fail to reject the null hypothesis for), we avoid placing a read if it has too few k-mer matches 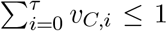 default: τ=2).

### Computing Parent and Abelian group hashing

The encodings of colors in **C** play an important role in the efficient computation of Parent(**G**, C_1_, C_2_). A desired encoding should enable fast computation of C = C_1_ ∪ C_2_, and we also need to query if C was already added to the multitree, G. We do this by using an Abelian group hashing scheme for sets, i.e., colors.

Using a hash function H with a sufficiently large range, we assign a hash value to each singleton color H({r}) r ∈ ℛ, . Then, it is possible to form an Abelian group by setting non-singleton colors’ hash values to H(C) = _*r∈C*_ H({r}). Since we only compute Parent of disjoint colors, we could obtain H(C) by summing H(C_1_) + H(C_1_). Thus, **G** could be stored in an associative array where we keep H(C) as the key and either H(C_1_) or H(C_2_) as the value, since one can always compute the other by subtraction. When C ∉ **G**, we simply add it, and return H(C_1_) + H(C_2_). When C ∈ **G**, it could be that it is either already seen due to another k-mer or it is a collision. We check collisions by comparing the value associated with H(C) and H(C_1_), H(C_2_). Despite being extremely rare, collisions are possible. We handle collisions by adding a dummy reference, i.e., a nonce, to the color C. At the beginning, we ensure that there is no collision among colors corresponding to internal nodes (and singletons) by rehashing.

At the end, in order to avoid storing redundantly large hash values, we simply convert colors to integers by enumerating them (using log(|𝒞|) bits) and store them in an array where indexes correspond to color encodings, instead of an associative array.

## Derivative of the likelihood function

Given, u_*r*_, **v**_*r*_, δ, k, h and ρ_*r*_, we can compute the derivate of the log-likelihood, Eq. (2), with respect to D as follows:

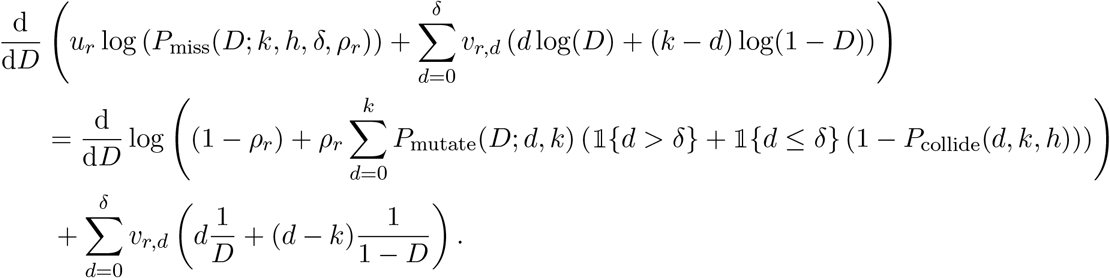

For convenience, we define the following function P_*δ*_(d): [0, k] → [0, 1],

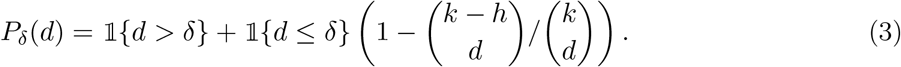

where 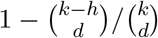 is simply 1 − P_collide_(d, k, h).

Then, we can substitute P_mutate_(D; d, k) with 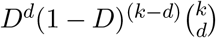 and compute the derivative of the logarithm,

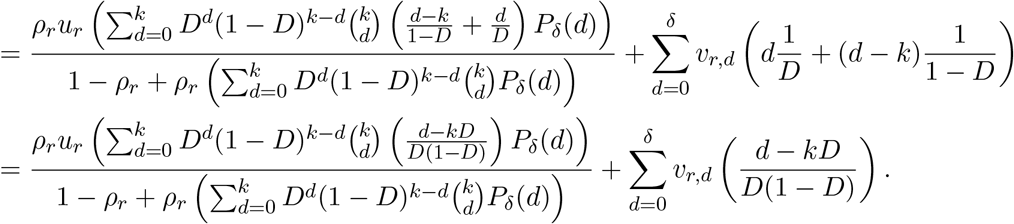

The sign of this derivative is positive at 0 + ϵ, and negative at 1 − ϵ, which is required for Brent’s method to work and find local optima.

The second derivative is given by

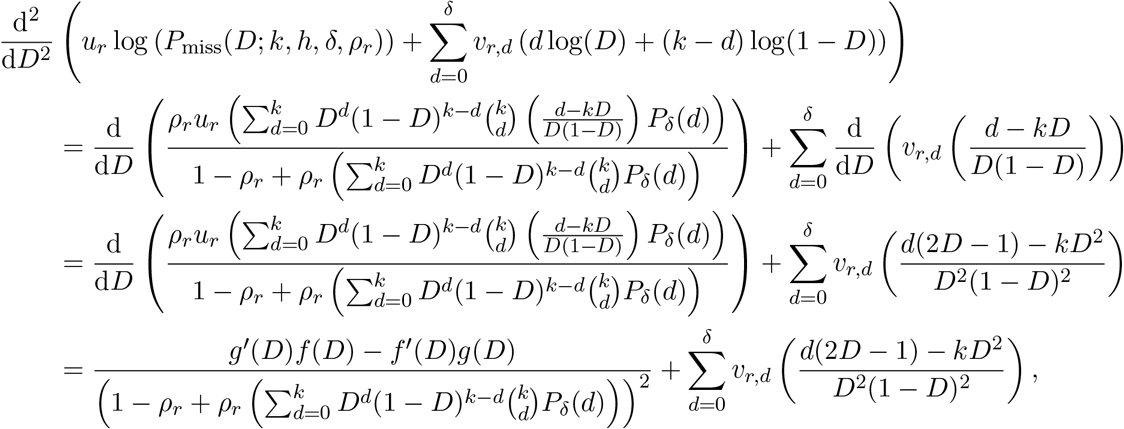

where

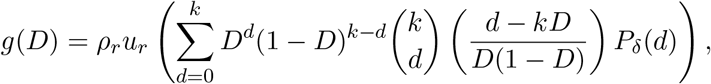

and

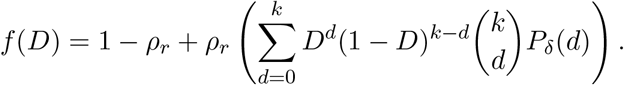

Here, the denominator of the first term is always positive, and the second term, due to P_match_ is non-positive for D ∈ (0, 0.5). Derivatives of f(D) and g(D) are given by

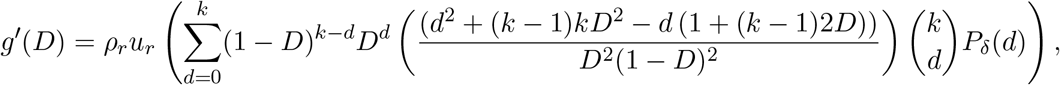

and

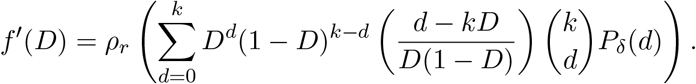

Thus, f ^*′*^(D)g(D) is always non-negative:

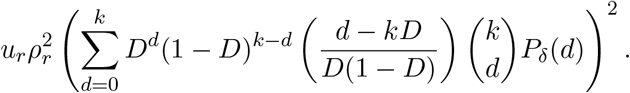

It suffices to show g^*′*^(D)f(D) ≤ 0 to prove the concavity of the log-likelihood. Notice that we can simplify the entire second derivative without changing the sign by multiplying all terms with D^2^(1 D)^2^. Furthermore, since f(D) > 0 for ρ_*r*_ ∈ [0, 1] and D (0, 0.5), we can focus on the following quantity:

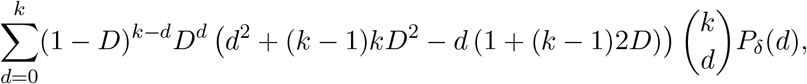

which is equal to 0 without P_*δ*_(d):

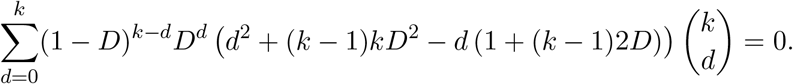

This can be shown by

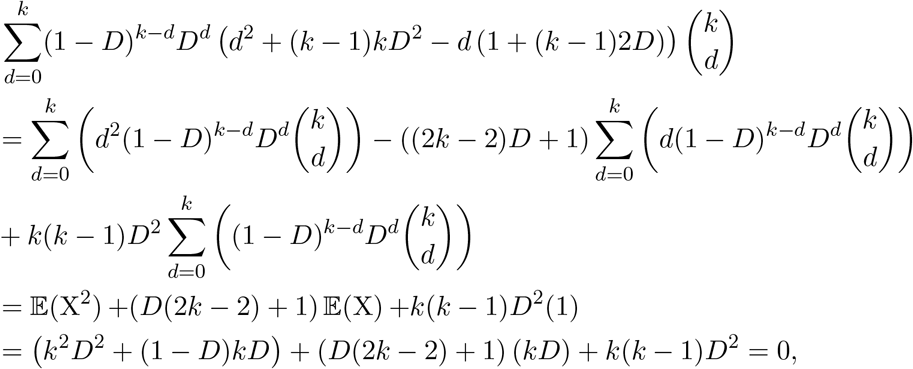

where X ∼ Binomial(k, D).

Finally, to conclude that g^*′*^(D) ≤ 0, we need to show the following inequality holds after including P_*δ*_(d) back:

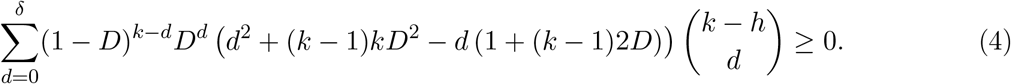

This inequality is not true for all choices of δ, h and k, but the default parameter values k=29, h=14, δ=4 satisfy it for D ∈ (0, 0.5)

### Why not APPLES?

Reads come from various parts of the genome, and the rates of evolution and, thus, branch lengths change across the genome. For the least square method APPLES [53] to work well, distances from query to references should be in the same scale as distances among references. This observation was one of the main insights from the original papers. Having consistent estimates of branch length necessitates recomputing distances among references for each read, which is possible with our data structure. However, it will make placement too slow as it would need O(kn^2^) computations for k queries and n references, instead of the desired O(kn), provided by krepp. Ignoring changes in rates reduces accuracy and is not a viable option [63].

### Software versions and commands used

Here, we provide the exact commands that we used to run external tools and krepp throughout our experiments, together with their version information.

#### Genomic distance estimation using Mash

We used Mash (version 2.3) to estimate genomic distances. To create a Mash sketch from a genome and then use it to estimate genomic distance, we used the below commands.

~~~
mash sketch -k 29 -s 100000 -p $NUM_THREADS -o $SKETCH_FILE $INPUT_FASTA
mash dist $SKETCH_FILE1 $SKETCH_FILE2
~~~

#### Short read simulation using ART

We simulated short reads with length L and coverage c, with the default error and quality profiles of Illumina HiSeq 2500 using ART [65] (version 2.5.8 – single read mode) with the command below.

~~~
art_illumina -ss HS25 -l L -f c -na -s 10 -i $INPUT_FASTA -o $OUTPUT_FASTQ
~~~

#### Downsampling reads using seqtk

To subsample read collection down to a specified number of reads, denoted by n here, we used `seqtk` [66] (version 1.3r106) with the command below.

~~~
seqtk sample -s 150 $INPUT_FASTQ n > $OUTPUT_FASTQ
~~~

#### Read alignment using bowtie2

We used bowtie2 [22] (v2.4.1) for short read alignment. To construct an index from all genomes combined in a single FASTA file (`$INPUT_FASTA`), we used the command below.

~~~
bowtie2-build --large-index --threads 32 $INPUT_FASTA $OUTPUT_DATABASE
~~~

Given an index ($INPUT_DATABASE), we used the following alignment configuration to generate SAM files for all alignments.

~~~
bowtie2 -p $NUM_THREADS -x $INPUT_DATABASE -t -q -U $QUERY_FASTQ \--xeq --very-sensitive --all -S $OUTPUT_SAMFILE
~~~

We postprocessed `$OUTPUT`_SAMFILE to compute Hamming distance for each alignment using a custom script provided at https://github.com/bo1929/shared.krepp/blob/main/scripts/postprocess_sam.py.

#### Distance estimation and phylogenetic placement using krepp

The version of krepp we presented in this work is v0.4.5. Given a mapping between reference IDs (leaves of the tree) and paths, together with an optional guide tree, krepp builds an index from k-mers of the reference genomes in batches, and the total number of batches is specified by the option -m. Each batch is built with a separate command using the option -r as shown in the below command.

~~~
krepp index -o $OUTPUT_INDEX -i $INPUT_FILE -t $BACKBONE_NEWICK --no-frac \
--num-threads $NUM_THREADS -k k -w w -h h-m $NUM_BATCHES -r $BATCH_INDEX
~~~

For WoL indexes, we set k = 29, w = 35, and h = 14. For larger RefSeq snapshots, we opt for k = 30, w = 37, and h = 14.

Once the index is built, one can query reads against it for either distance estimation or phylogenetic placement using the following commands, setting δ = 4:

~~~
krepp dist -i $INPUT_INDEX -q $QUERY_FASTQ -hdist-th δ \
--num-threads $NUM_THREADS -o $OUTPUT_DISTANCES
krepp place -i $INPUT_INDEX -q $QUERY_FASTQ –hdist-th δ \
--num-threads $NUM_THREADS -o $OUTPUT_JPLACE
~~~

#### BIOM table construction using Woltka

We converted mappings of short reads to BIOM tables, which are simply matrices of counts of observations on a per-sample basis, using Woltka (version 0.1.7). In particular, we ran `woltka classify --no-demux --digits 5 -i $INPUT_MAPPINGS -o $OUTPUT_BIOM`, where $INPUT_MAPPINGS is a directory containing tab-separated files for read ID to subject (e.g., reference genome, internal node, taxon) mappings for each sample. To filter low-abundance subjects, we used `woltka filter -i $INPUT_BIOM -o $OUTPUT_BIOM --min-percent 0.01`.

#### Genome-wide phylogenetic placement using App-SpaM

We used App-SpaM [11] (v1.03) to place short reads with respect to reference sequences, combined in a single FASTA file (`$INPUT_FASTA`), on a given backbone tree (`$BACKBONE_NEWICK`) by running the command below.

~~~
appspam --threads $NUM_THREADS -s $INPUT_FASTA -t $BACKBONE_NEWICK -q $QUERY_FASTA
~~~

#### Marker-based phylogenetic placement using EPA-ng

For maximum likelihood placement, the branch lengths of the backbone tree are re-estimated using RAxML [81] (v1.2.2) using the command below.

~~~
raxml-ng -evaluate --threads $NUM_THREADS --msa $REFERENCE_MSA \
--tree $BACKBONE_NEWICK --prefix $OUTPUT_PATH/mltree \
--model GTR+G+F --force --blopt nr_safe
~~~

Using the output tree from the above command, we used EPA-ng (version 0.3.8) as shown below to place queries

~~~
epa-ng --ref-msa $REFERENCE_MSA --tree $BACKBONE_NEWICK --query $QUERY_FASTA \
--model $OUTPUT_PATH/mltree.raxml.bestModel -w $OUTPUT_DIR \
--verbose --redo -T $NUM_THREADS
~~~

#### Rarefication and psedo-F calculation using QIIME2

We computed separation statistics and distances between sample pairs using QIIME2 [69] (version 2024.5.0), and specifically its diversity plugin (version 2024.5.1). We utilized `core-metrics-phylogenetic` command to compute weighted UniFrac distance and Bray-Curtis dissimilarity after performing rarefication at sampling depth S (set to 100, 000 and 6, 550 for human microbiome and Earth’s microbiome analyses, respectively).

~~~
qiime diversity core-metrics-phylogenetic --i-table $FEATURE_TABLE_QZA \
--i-phylogeny $BACKBONE_QZA --m-metadata-file $METADATA_FILE \
--p-sampling-depth S --p-ignore-missing-samples --output-dir $OUTPUT_DIR
~~~

Next, once the distance matrices are obtained, we used beta-group-significance command to compute pseudo-F statistics for all groupings as shown below.

~~~
qiime diversity beta-group-significance --i-distance-matrix $DISTANCE_MATRIX_QZA \
--m-metadata-file $METADATA_FILE --m-metadata-column $METADATA_COLUMN \
--o-visualization $OUTPUT_QZV --p-permutations 1000
~~~

